# Division state reveals hidden genetic regulation during T cell activation and identifies immune disease–linked gene programmes

**DOI:** 10.64898/2026.02.22.707277

**Authors:** Megan Gozzard, Kyuto Sonehara, Kevin Ly, Tarran Rupall, Ziying Ke, Ximena Ibarra-Soria, Mohammad Lotfollahi, Blagoje Soskic, Carla P. Jones, Olivier B. Bakker, Gosia Trynka

**Affiliations:** Wellcome Sanger Institute, Wellcome Genome Campus, Hinxton, CB10 1RQ, UK; Cambridge Center for AI in Medicine, University of Cambridge, Cambridge, UK; Research Technologies, GSK, Stevenage, UK; Cambridge Stem Cell Institute, University of Cambridge, Cambridge, UK; Human Technopole, Viale Rita Levi-Montalcini 1, 20157, Milan, Italy; Open Targets, Wellcome Genome Campus, Hinxton, CB10 1RQ, UK

## Abstract

A major challenge in human genetics is to connect disease-associated variants to the cellular functions they perturb. Variants associated with immune-mediated disease have implicated CD4⁺ T-cell activation as a disease-causal process; however, it remains unresolved whether these variants act on pathways initiating cell division, driving effector differentiation, or maintaining proliferative capacity. Activated T cells comprise mixtures of cells that have undergone different numbers of divisions, potentially obscuring genetic effects that act at specific stages of this process. Here, we study CD4⁺ T-cell activation through cell division using division-resolved single-cell transcriptomics of naive, memory and regulatory T cells. We show that division state constitutes an important axis of transcriptional organisation, with approximately one third of genes exhibiting division-dependent regulation across shared and subset-specific programmes. We develop *CellDivider*, a machine-learning model that accurately infers proliferative state directly from single-cell transcriptomes, enabling a scalable approach for cohort level mapping of variant effects on cell proliferation phenotypes. Incorporating inferred division state into single-cell eQTL models reveals regulatory effects that are attenuated or missed when cells are analysed by activation time alone. Finally, we resolve gene expression programmes that govern distinct stages of T cell activation and proliferation. We show that polygenic immune disease risk preferentially converges on early activation and differentiation modules, rather than on proliferative capacity itself. Consistent with this model, division-resolved analysis of the pleiotropic immune risk gene *RPS26* localises its genetic effects to early division states, providing mechanistic insight into its broad disease associations. Together, these findings illustrate how resolving dynamic cellular consequences of activation sharpens variant-to-function inference in immune disease genetics.

## Introduction

Over the past decade, the transition in complex disease genetics research towards single cell resolution has sparked a growing recognition that the functional effects of GWAS variants are highly context dependent ^1^. While early studies established that genetic risk manifests in a cell type–specific manner ^2^, it has become clear that cell type alone is insufficient to explain where and how disease-associated variants exert their effects ^3,4^. Many regulatory variants act only in particular cellular states, such as during activation, differentiation, or stress responses, when relevant regulatory elements become accessible and gene regulatory programmes engaged ^5^. Failure to account for this contextual variability may confound the interpretation of genetic effects, analogous to the averaging gene expression inherent in bulk RNA sequencing, which integrates signals across heterogeneous cell populations and thereby obscures cell-type–specific regulatory mechanisms ^6^. In single-cell studies, genetic effects can either remain hidden, or lead to confounded interpretation, if contextual heterogeneity within a cell type is not explicitly modelled ^7,8^. The extent of regulatory variants which are influenced by context is substantial. For instance, expression quantitative trait loci (eQTL) mapping in resting and activated CD4+ T cells has revealed highly dynamic gene expression regulation. Of 6,407 cis-eQTLs identified during CD4+ T cell activation, 35% showed a change of effect size over time, with ∼22% of eGenes detected exclusively at the proliferative phase, five days post-activation ^9^. These findings suggest that many genetic effects are conditional not simply on activation, but on specific stages of the T cell response.

Therefore, disentangling these layers of context is essential for resolving both individual variant effects and polygenic signals, as well as for linking genetic risk to the specific cellular phenotypes that are dysregulated in disease. However, defining context remains challenging. For most cell types, we lack a clear understanding of which cellular states and phenotypes are most relevant to disease, and the space of possible contexts, e.g. spanning activation, differentiation, proliferation and other functional trajectories, is vast.

GWAS in immune-mediated disease (IMD) have established a causal role for CD4⁺ T cells in disease pathogenesis, with risk variants enriched in regulatory elements active in these cells ^10–13^. The enrichment is particularly strong in enhancers and promoters that become upregulated during T cell activation compared with the resting state ^10^. However, given the breadth and complexity of the contextual landscape CD4+ T cells inhabit, previous SNP enrichment analyses in immune disease have implicated broad cellular states like ‘T cell activation’ rather than the specific cellular processes governing disease risk. In part this is due to CD4^+^ T cell activation occurring over several phases with the key stages being early activation (up to 40h after effective stimulation, considered here as the regulation that governs the cells decision to divide) ^14^ versus late activation, which is where CD4+ T cells undergo rapid clonal expansion and they acquire and refine their effector functions. T cell proliferation is both a hallmark of protective immunity and a driver of chronic inflammation in immune-mediated disease. Upon antigen recognition and activation, naive CD4+ T cells initiate a tightly orchestrated programme of expansion, to the extent that CD4+ T cells can proliferate up to 5,000-fold ^15^. Following the proliferation phase, most cells undergo apoptosis, while a subset remodels their epigenetic and transcriptional landscapes to become memory T cells with faster recall capacity ^16,17^. Notably, CD4⁺ T cell abundance itself is genetically regulated, it varies across the lifespan and between sexes, and shares genetic associations with immune disease susceptibility ^18–20, 21^. As such, T cell proliferation is a direct outcome of cell stimulation and a defining hallmark of cell activation. However to what extent cell proliferation, or other activation-linked cell phenotypes, play a causal role in complex immune disease remains to be determined.

To address this gap, we performed *in vitro* profiling of cell division in naive, memory, and regulatory CD4⁺ T cells, the major CD4⁺ T-cell subsets with distinct activation dynamics and effector functions, across successive rounds of division from the first to the fifth. We generated a single-cell transcriptional atlas of 112,454 cells and identified division-dependent gene expression changes. We trained a multilayer perceptron model, *CellDivider*, and applied it to transcriptomic data from a CD4⁺ T-cell activation time course ^9^ which revealed substantial inter-individual heterogeneity in proliferative outcomes. Incorporating this inferred division state into single-cell interaction eQTL models uncovered additional layers of gene regulation that were not captured when cells were grouped solely by activation timepoint. Division state provided a new context for interpreting regulatory effects that enabled interpretation beyond coarse activation stages.

Finally, we resolved gene-expression programmes that capture the molecular transitions accompanying proliferation, spanning cell-cycle entry, metabolic adaptation, cytokine induction, and effector acquisition. Integrating these programmes with immune-disease genetics identified polygenic IMD risk preferentially enriched in specific transcriptional modules.

## Results

### Cell division uncovers dynamic and subpopulation-specific transcriptional regulation

To map the transcriptomic programmes underlying CD4⁺ T cell proliferation, we isolated three major CD4⁺ T cell populations: naive, memory, and regulatory T cells (Tregs), from four healthy donors (**Methods**). Cells were labelled with CellTrace Violet (CTV) and stimulated with aCD3/aCD28/aCD2 (Immunocult). Four days after stimulation, CD4⁺ T cell cultures comprised a heterogeneous mixture of cells that had undergone different numbers of divisions. We sorted cells based on CTV intensity, which halves with each successive cell division, allowing us to capture six distinct division states. Each sorted fraction (referred to as a division state) included an equal number of cells and was subject to single-cell transcriptomic profiling (**Fig. 1a** and **Methods**). The final dataset comprised 86,760 high-quality single cells, with 10,058–15,536 cells per division state.

**Figure 1.**
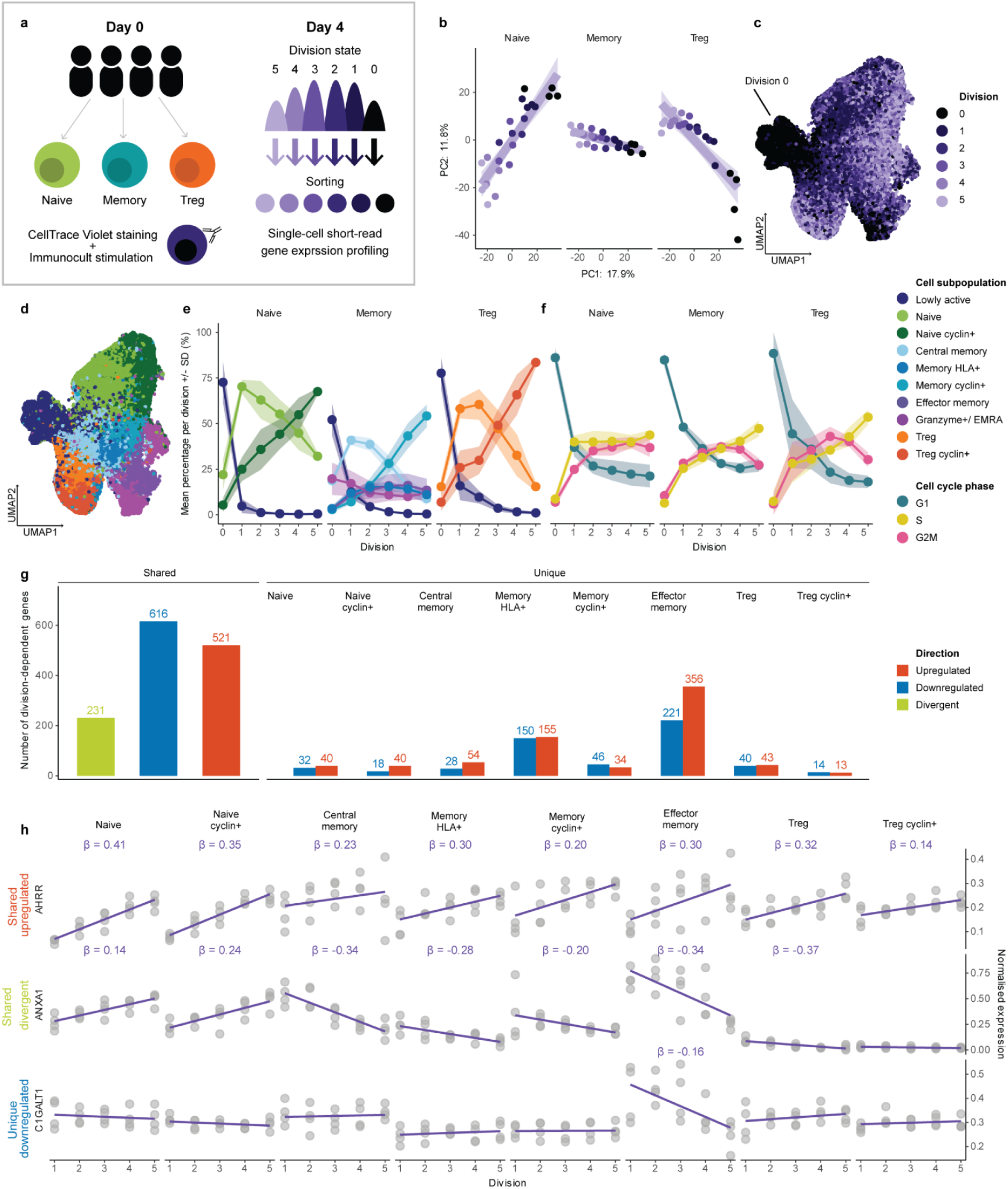
Division-dependent gene expression reveals shared and divergent activation signatures. **a.** Overview of the study design. Briefly, naive, memory and Treg cells were isolated, labelled with cell trace violet (CTV) and stimulated with Immunocult. Four days after stimulation cells were sorted based on the intensity of CTV, corresponding to the number of divisions a cell had undergone. Sorted cells from each cell population and across all divisions were transcriptionally profiled with single cell resolution. **b.** Principal component analysis of aggregated expression across population–division–donor, coloured by division state. Percentages at the axes depict gene expression variance explained by principle components (PC) 1 and PC2. **c-d** UMAP embedding of 86,760 cells in the final dataset, coloured by division state **(c)**, and transcriptionally-defined subpopulation (subpopulation key shown in figure legend) **(d)**. **e.** Proportion of subpopulations across division states and cell populations (mean +/− standard deviation). **f.** Proportion of cells in inferred cell cycle phases across division states and cell populations (mean +/− standard deviation). **g.** Division-dependent differentially expressed genes, grouped as divergent (significantly different across subpopulations), shared (significant in more than one subpopulation) or unique (restricted to a single subpopulation) (p_adj_ < 0.05, |β| > 0.1). **h.** Example genes with different division-dependent effects. Plots depict normalised gene expression : *AHRR* (globally upregulated), *ANXA1* (divergent across subpopulations), and *CGALT1* (specifically downregulated in effector memory cells). β values shown where p_adj_< 0.05, |β| > 0.1.

Dimensional reduction analysis revealed a clear transcriptional gradient corresponding to the number of cell divisions (**Fig. 1b-c**) across all T cell populations. When evaluated per gene, division state explained up to 80% of pseudobulk and 49% of single-cell gene expression variance (mean 12.6% and 3.8%, respectively) (**Methods** and **Supplementary Fig. 1a-b**). Naive, memory, and Tregs clustered into 10 cell subpopulations (Louvain clustering resolution 0.2–0.5; **Fig. 1d**), which we labelled as naive, naive cyclin⁺, central memory, memory cyclin⁺, memory HLA⁺, effector memory, Granzyme^+^/EMRA, Treg, and Treg cyclin⁺ cells (**Fig. 1d, Supplementary Fig. 2a-c** and **Methods**). Additionally, lowly active cells ^9^ (**Fig. 1c-d**) composed of undivided cells from all populations clustered together, suggesting a distinct transcriptomic profile which was characterised by expression of *KLF2, IL7R, TCF7*.

Cluster composition shifted across division states: later divisions showed a high proportion of cells expressing mitotic cyclin genes (*CCNB1, CCNB2, CDC20, CDKN3*) (**Fig. 1e, Supplementary Fig. 2b**). These genes encode proteins essential for the G1/S transition and mitosis, consistent with the higher abundance of actively proliferating cells expected in later pools of divided cells. Cell cycle phase inference further revealed a progressive decrease in the proportion of cells in G1 phase with increasing division number, indicating that repeatedly dividing cells remain highly activated (**Fig. 1f**). Once cells entered division 2, the proportion of cells in each cell cycle phase remained relatively stable (**Fig. 1f**), indicating that the observed transcriptional heterogeneity across divisions is not solely explained by cell-cycle phase and is a distinct phenotype (**Supplementary Fig. 1c-d**).

Next, we examined if gene expression changes were associated with cell division and whether these effects were shared between cell subpopulations or subpopulation-specific (i.e. only significant in one subpopulation). Across the annotated subpopulations, of the 7,832 tested genes we identified 2,696 differentially expressed genes whose expression correlated linearly with division state in at least one cell subpopulation (p_adj_<0.05, |β| > 0.1) (**Fig. 1g, Supplementary Table 1**). Using gene set enrichment analysis (GSEA) and the Hallmark gene set collection (*h.all.v2025.1.Hs*) division-dependent genes revealed distinct enrichments (FDR < 0.05) across CD4⁺ T-cell subpopulations, reflecting division state-specific functional remodelling during proliferation. We observed significant enrichment for metabolic and mitochondrial pathways in division-dependent upregulated transcripts, including mTORC1 signalling and oxidative phosphorylation, key regulators of T-cell metabolism and proliferative capacity. Conversely, immune-related pathways such as IFN-α and IFN-γ response, TNF-α and TGF-β signalling, and the inflammatory response were broadly downregulated, consistent with the attenuation of early activation signals as cells transition into proliferative phases (**Supplementary Fig. 3**).

Over half of the 2,696 division-dependent genes (52%, n =1,412) were shared across two or more subpopulations, including 521 consistently upregulated and 616 consistently downregulated with division (**Fig. 1g**), e.g., *AHRR*, which encodes the aryl hydrocarbon receptor repressor that regulates CD4⁺ T-cell differentiation and function, was consistently upregulated across all subpopulations (**Fig. 1h**). We asked whether the magnitude or the direction of division-dependent transcriptional responses differed significantly between CD4⁺ T-cell subpopulations. Divergent division-dependent effects were modelled using an interaction framework in which gene expression was regressed on division number with cell subpopulation–specific interaction terms, testing whether the transcriptional response to successive rounds of division differed between CD4⁺ T-cell subpopulations relative to naive cells. We identified 231 unique genes with divergent effects, corresponding to 302 significant gene-division interaction coefficients (p_adj_ < 0.05) (**Fig. 1g, Supplementary Table 2-3**). The majority of divergent coefficients were negative (230/302, ∼76%), indicating that relative to the naive cells, gene expression of these genes is repressed as memory and Treg subpopulations progress through division. This could indicate that memory and Treg populations are specializing their transcriptome to support their effector functions as they divide, whereas naive cells retain more plasticity in their transcriptome to enable differentiation.

To assess if these genes were involved in specific processes, we applied GSEA using GO Biological Processes and the Hallmark gene set collections (*c5.go.bp.v2025.1.Hs*, *h.all.v2025.1.Hs*; FDR < 0.05) showed that genes attenuated in memory and Treg subsets were enriched for lymphocyte activation (e.g. *CD40LG, LCP2, LAT*), cell–cell adhesion (e.g. *CD58, ITGAL, ICAM3*), and cytokine, inflammatory and stress-response pathways, including JAK–STAT signalling (e.g. *STAT1, PIM1*), IL2–STAT5 signalling (e.g. *IL2RA, SOCS1*), TNF-α signalling (e.g. *TNFAIP3, TRAF1*), and the IFN-α and IFN-γ responses (e.g. *IFIT3, ISG15, OAS1*) (**Supplementary Fig. 4**). Divergent genes also included those involved in TCR signalling, co-stimulation and immune synapse organisation, including *CD40LG, CD58, CD48, CD226, CD82* and adaptor/scaffold molecules such as *ANXA1/2/5/6, UBASH3A, TRADD, TRAF1* and *TNIP3*. Notably, *ANXA1* showed the strongest opposing division-associated trends, increasing with division in naive cells but decreasing in memory populations, particularly effector memory cells (**Fig. 1h**). This pattern is in line with the role for *ANXA1* in tuning activation thresholds during initial naive-cell activation, while its downregulation in antigen-experienced cells may serve to constrain proliferative or inflammatory responses once activation history has been established ^22,23^.

Finally, we also observed extensive subpopulation-specific regulation: 1,284 genes were division-dependent in only one subpopulation (735 upregulated, 549 downregulated, (p_adj_ < 0.05, |β| > 0.1; **Fig. 1g, Supplementary Table 3**). One example, with a strong effect size manifesting specifically in effector memory cells (βEffector memory = −0.16 in comparison to β in other subpopulations (−0.017 to 0.04, p_adj_ >0.23), is *C1GALT1,* which encodes a key enzyme that controls the O-glycosylation of surface proteins critical for trafficking. Although *C1GALT1* expression was generally low and stable across division states and CD4⁺ T-cell subpopulations (mean normalised expression 0.25–0.34), effector memory cells showed a distinct division-dependent pattern. In this subset, *C1GALT1* expression was relatively elevated in early divisions (mean normalised expression_Division_1_ = 0.42) and progressively declined with successive divisions, reaching levels comparable to other subpopulations by Division 5 (mean normalised expression_Division_5_ = 0.25) (**Fig. 1h**). This early, transient upregulation may reflect a short-lived requirement for robust core 2 O-glycan synthesis, to support tissue homing ^24,25^.

Collectively, these findings demonstrate that division state is associated with structured transcriptional programmes modulated by activation history. We show that division state is a significant axis of gene expression variation, encompassing coordinated metabolic, signalling, and effector pathways, encoded in transcriptomic signatures.

### *CellDivider:* a transcriptome-based model to accurately predict proliferative state of activated T cells

Measuring proliferation at high throughput is challenging because it relies on bespoke biological assays. It requires isolating live cells, labelling them with division dyes, and quantifying successive divisions by flow cytometry. Scaling such assays to hundreds of individuals is difficult, technically demanding, and requires specialised expertise. Based on the substantial transcriptional variability explained by division state, we hypothesised that our dataset of division-resolved single-cell transcriptomic profiles in naive and memory CD4⁺ T cells contains gene expression signatures sufficient to learn division state directly from transcriptomes, enabling population-scale modelling of proliferation.

To test this, we developed a machine learning model, *CellDivider*, to predict the division state of each cell based on transcriptomic data (**Fig. 2a** and **Methods**). We tested multiple architectures (elastic-net regression, eXtreme Gradient Boosting and multi-layer perceptron, MLP) and normalization strategies with the hyperparameters optimised by a five-fold cross-validation framework. To assess the generalisability of these models, we generated an independent single-cell transcriptomic test dataset comprising 25,694 activated CD4⁺ T cells also sorted by CTV-labeled division state at 3 days and 5 days after activation (**Fig. 2b** and **Methods**). We opted to predict division states as a continuous rather than a discrete outcome to better capture the inherent uncertainty in the predictions, while the experimentally observed division states are always discrete by definition (**Methods**). When applied to the test dataset with both timepoints, the MLP model using scVI-adjusted gene expression as input features achieved the highest predictive accuracy (Pearson *r* = 0.881; Root Mean Squared Error [RMSE] = 0.705; Mean Absolute Error [MAE] = 0.428; **Fig. 2c; Supplementary Fig. 5**). Division-dependent differential gene expression was highly correlated between Day 3 and Day 5 (R= 0.83, p < 2.2 × 10⁻^16^; **Supplementary Fig. 6**), indicating that the underlying molecular features of proliferation are largely time-independent.

**Figure 2.**
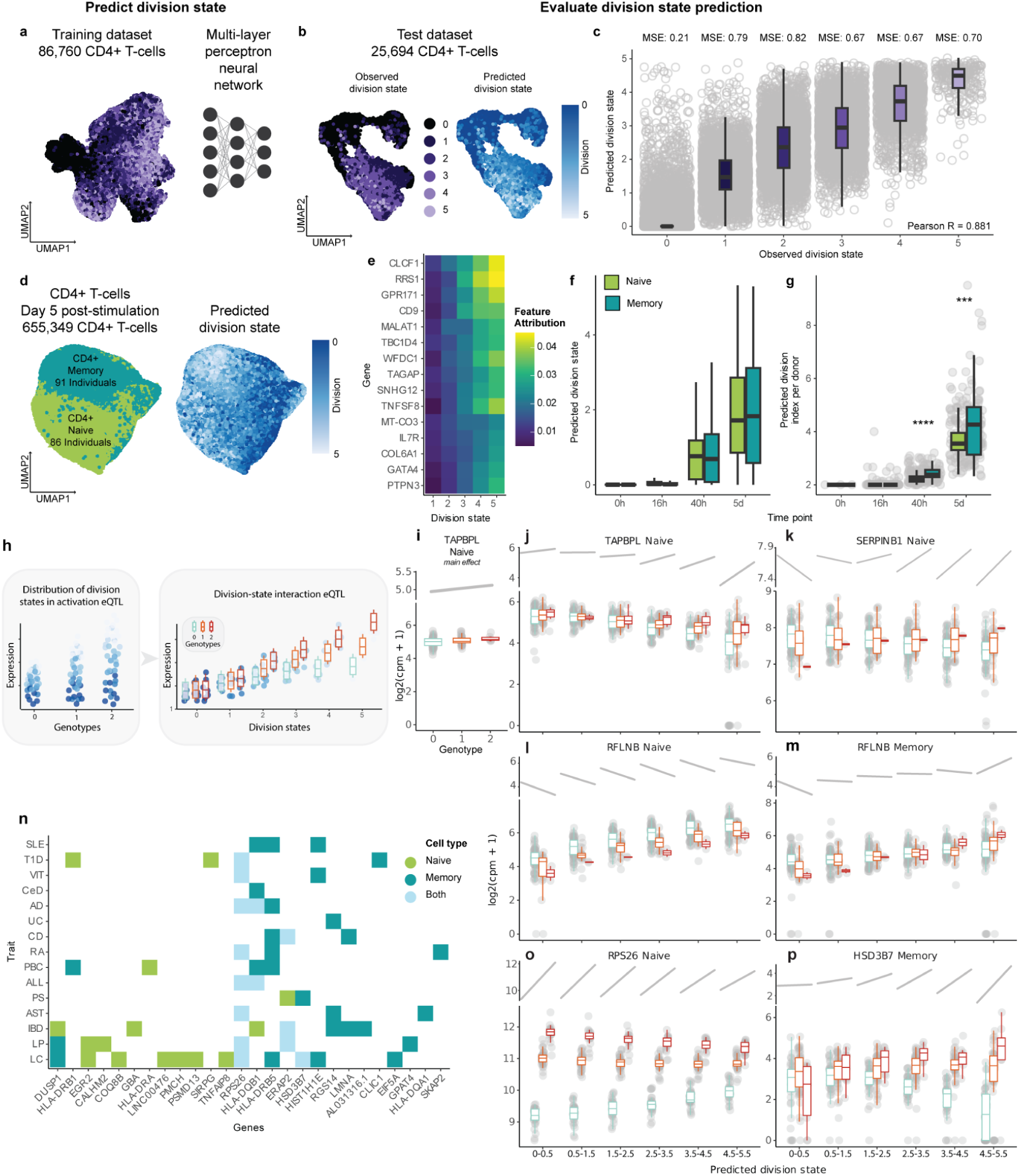
Transcriptome-based inference of T cell proliferation resolves context-dependent eQTLs. **a.** Schematic of the development of the *CellDivider* model that predicts division state from single-cell transcriptome. **b.** UMAP embedding of an independent test dataset of 25,694 CD4+ T cells. The cells are coloured by experimentally observed division state in the left panel (purple) and by computationally predicted division state in the right panel (blue). **c.** Distribution of the predicted division state, stratified by the experimentally observed one. Each point represents a cell. MSE, mean squared error. **d.** UMAP embedding of a population-scale dataset of 655,349 post-stimulation CD4+ T cells. The cells are coloured by cell population (CD4+ naive/memory) in the left panel and predicted division state (blue) in the right panel. **e.** Heatmap showing feature attribution computed using integrated gradients. **f-g.** Distribution of predicted division state per cell **(f)** and division index per donor calculated from predicted division state **(g)**. Significance levels are indicated as *** (p <= 0.001), **** (p <= 0.0001). **h.** Schematic of the division-state-dependent eQTL analysis. The genotype×division state interaction effect was evaluated using a single-cell eQTL modelling framework. **i.** Genotype–gene expression associations visualised for the G allele at chr12:6479279 and *TAPBPL* in naive cells modelled for the main effect (p = 0.18), each point represents the average log_2_(CPM + 1) across all cells in a donor, grouped by genotype. The top sloped line is based on a linear regression of log_2_(CPM + 1) against genotype. **j-m.** Examples of division-state-dependent eQTL. Genotype–gene expression associations visualised for the G allele at chr12:6479279 and *TAPBPL* in naive cells at the division-state-dependent effect (p_interaction_ = 3.6 × 10^−6^) **(j)**, the C allele at chr6:2823093 and *SERPINB1* in naive **(k)**, the A allele at chr17:525981 and *RFLNB* in naive **(l)** and memory **(m)**. **n.** Tile plot showing eGenes with significant division-state-dependent eQTL effects by GWAS lead variants. Colours indicate the cell population in which the eQTL effect was observed. GWAS abbreviations: LC, lymphocyte count; LP, lymphocyte percentage; IBD, Inflammatory Bowel Disease; AST, Asthma; PS, Psoriasis; ALL, Allergic Disease, PBC, Primary Biliary Cholangitis; RA, Rheumatoid Arthritis; CD, Crohn’s Disease; UC, Ulcerative Colitis; AD, Atopic Dermatitis; CeD, Coeliac Disease; VIT, Vitiligo; T1D, Type 1 Diabetes; SLE, Systemic Lupus Erythematosus. In all panels, boxplots represent the interquartile range (IQR), ends of whiskers represent minimum and maximum values within 1.5 × IQR. **o-p.** Genotype–gene expression associations visualised for the C allele at chr12:56042145 and *RPS26* in naive **(o)** and the G allele at chr16:30999862 and *HSD3B7* in memory **(p)**. Panels **j-m, o and p**, each point represents the average log_2_(CPM + 1) across all cells in the indicated division state bin in a donor, grouped by genotype. The top sloped lines are based on a linear regression of log_2_(CPM + 1) against genotype in each division state bin. CPM, counts per million.

Having established that our predictive model was highly accurate and applicable at various post-activation timepoints, we applied *CellDivider* to a previously published single-cell dataset of 655,349 naive and memory CD4⁺ T cells collected from 119 healthy individuals at rest (0h) and after three activation time points (16h, 40h and 5 days) (**Fig. 2d**) ^9^. We generated scVI-adjusted gene expression profiles derived by jointly integrating and batch-correcting our main, test, and *Soskic et al.* datasets. Using this merged dataset, we trained *CellDivider* on cells from the main dataset and evaluated its performance on cells from the held-out test dataset. We observed a similarly high predictive accuracy (Pearson *r* = 0.877; RMSE = 0.676; MAE = 0.409). Genes contributing most strongly to model performance included *CLCF1, RRS1, GPR171, CD9* and *MALAT1* (**Fig. 2e, Supplementary Table 4**). This enabled us to assign cells to continuous predicted division states ranging from zero to five (**Fig. 2f-g**, **Supplementary Fig. 7**).

At 0h and 16h no cells were predicted to have undergone division but at 40h after stimulation, a time window during which CD4⁺ T cells are expected to have completed at least one division, we observed the majority of predicted division states for both naive and memory subsets within a 0–1 range (**Fig. 2f, Supplementary Fig. 7**). At five days, the median predicted division state differed slightly between cell populations, with a median 1.71 in naive cells and 1.82 in memory cells. These observations build our confidence in the accuracy of *CellDivider’s* modelling.

Next we assessed the inter-individual variability in predicted division states by calculating the proliferation index (the average number of cell divisions undergone by all cells in the original starting population, including those that never divided), division index (the average number of divisions undergone only by the cells that have divided at least once) and the percent divided (the percentage of initial cells that underwent at least one division) for each donor, time point and cell population. We observed substantial inter-individual variability in division indices, with variability more pronounced in memory than in naive populations, and dividing memory cells expanding more compared to naive cells (**Fig. 2g**, **Supplementary Fig. 8**, mean division index_Naive_ = 3.7, mean division index_Memory_ = 4.3, p = 5.3 × 10⁻^4^ at Day 5).

Given the inter-individual variability, we tested whether genetic variants shape the proliferative response of CD4⁺ T cells to activation, by performing GWAS on the division index, proliferation index and percent divided. While no variants reached genome-wide significance (p < 5 × 10⁻⁸) at the current sample size, we identified 292 suggestive association signals (p < 1 × 10⁻⁵) across CD4⁺ T cell proliferation traits in both naive and memory subsets (**Supplementary Table 5**), pointing to potential genetic contributors to proliferative variability. For instance, a signal within the *RPTOR* gene (p = 2.2×10⁻^6^, β = 0.68), the defining scaffold subunit of mTORC1, critical for licensing entry into proliferation. The absence of genome-wide significant signals is consistent with expectations for GWAS of complex cellular phenotypes which typically require hundreds to thousands of samples to achieve genome-wide significance. Nonetheless, even with a limited sample size, we identified a number of suggestive loci that may contribute to variation in cellular phenotypes.

Together, these results demonstrate that transcriptional signatures of proliferation are conserved, temporally stable, and quantitatively predictive of a cell’s division state. The *CellDivider* model provides a general framework to infer proliferative states from a transcriptomic dataset of activated CD4+ T cells, enabling systematic assessment of division-linked genetic effects without the need for experimental cell labelling.

### Division state uncovers context-dependent eQTL effects

Having established that division state can be accurately inferred from transcriptomic data and represents a major source of cellular heterogeneity during CD4⁺ T cell activation, we next asked whether variation in division state constitutes a biologically meaningful context for genetic regulation of gene expression (**Fig. 2h**).

We tested for division-dependent eQTLs at the single cell level through negative binomial mixed-effects (NBME). We first examined the distribution of association statistics using permuted datasets by randomly selecting eQTLs identified in the original *Soskic et al.* study and generated permuted datasets by shuffling donor–genotype assignments. We employed the recently proposed NBME model, which includes donor- and batch-specific intercepts as random effects (^8,9,26–28^; **Methods**). When we tested the main genotype effects in 1,000 permuted datasets using this model, the p-values obtained from the likelihood ratio test were uniformly distributed (λ, defined as 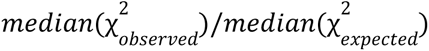, ranges 0.98–1.24), indicating that the statistical test was well-calibrated (**Supplementary Fig. 9a**). We next tested the interaction term between genotype and predicted division state using the same procedure. In contrast to the main effect, we observed systematic inflation of the association statistics for the interaction effect in the permuted datasets (λ ranges 0.95–1.95; **Supplementary Fig. 9b**). We reasoned that this inflation was caused by between-donor heterogeneity in the effect size of cellular state being erroneously attributed to genotype. To prevent this misattribution, we extended the NBME model by adding a donor-level random slope for the cellular state effect and repeated the permutation analysis (**Methods**). This extended model substantially improved calibration (λ ranges 0.95–1.15; **Supplementary Fig. 10**), and we therefore adopted it for subsequent analyses.

For each gene–variant pair previously identified as a lead eQTL at any time point or cell population in *Soskic et al.* (13,519 pairs, spanning 6,407 genes, of which 4,974 manifesting in naive and 4,415 in memory cells), we tested for division-dependent genetic effects. Notably, 1,488 of the 6,407 genes (23%) showed division-dependent differential expression in our dataset, suggesting that the proliferative state represents a substantial source of transcriptional variation among eGenes, motivating division-state-aware interaction modelling. In naive T cells, 203 variant–gene pairs (corresponding to 116 genes, 2.3%) showed significant division interactions (FDR < 0.1). In memory cells, 526 pairs (285 genes, 6.5%) were significant (**Supplementary Tables 6-7**). We observed a comparable proportion of effects manifesting in early and late divisions. In naive cells, 85 effects (42%) manifested predominantly in early division states (*e.g.*, 0–1 division) and 118 in late division states (*e.g.*, 4–5 division), while in memory cells, 250 (47%) in early and 276 in late division states. As expected, main eQTL effects were generally stronger than interaction effects (**Supplementary Fig. 11**) but examples such as *TAPBPL*, where the main effect was weaker compared to the division-interaction eQTL (p_main_ = 0.18 and p_interaction_ = 3.6 × 10^−6^ in naive cells) indicate that there are also effects that can be masked when not accounting for division state (**Fig. 2i and 2j**). Furthermore, this analysis revealed qualitatively distinct regulatory patterns across proliferative trajectories, e.g. flipped allelic effects across division states as observed for *SERPINB1*; (p_interaction_ = 2.7 × 10^−12^; β_early_ = −0.08 and β_late_ = 0.24; **Fig. 2k**). Finally, we also observed division-induced cell population specific effects. The A allele at chr17:525981 decreased *RFLNB* expression, with the strongest effect observed in naive cells at early division states (β_early_ = −0.22; **Fig. 2l**). In memory cells, the same allele also reduced *RFLNB* expression at early division states with comparable effect size (β_early_ = −0.20), however, this effect was reversed in later division states, where the A allele was associated with increased *RFLNB* expression (β_late_ = 0.047; **Fig. 2m**).

To assess whether immune-disease risk variants could also regulate gene expression in the division-state-specific context, we performed analogous interaction tests for the 1,263 lead GWAS variants for 15 unique immune-mediated diseases ^29–46^ and 916 lead variants for lymphocyte counts or percentages ^47^ (**Supplementary Table 8**). In naive T cells, 25 variant–gene pairs (corresponding to 9 genes: *RPS26*, *ERAP2*, *SIRPG*, *GBA*, *DUSP1*, *HLA-DRA*, *HLA-DRB1*, *HLA-DRB5*, and *HLA-DQB1*) among the disease variants and 13 pairs (corresponding to 10 genes: *RPS26*, *ERAP2*, *HSD3B7*, *PMCH*, *PSMD13*, *CALHM2*, *TNFAIP8*, *LINC00476*, *COQ8B*, and *EGR2*) among the lymphocyte trait variants showed significant division interactions (FDR < 0.1; **Fig. 2n**). In memory T cells, 44 pairs (corresponding to 13 genes: *RPS26*, *ERAP2*, *RGS14*, *HSD3B7*, *SKAP2*, *CLIC1*, *LMNA*, *HIST1H1E*, *HLA-DQA1*, *HLA-DQB1*, *HLA-DRB1*, *HLA-DRB5*, and *AL031316.1*) among the disease variants and 10 pairs (corresponding to 8 genes: *RPS26*, *ERAP2*, *HSD3B7*, *DUSP1*, *EIF5A*, *GPAT4*, *HIST1H1E*, and *HLA-DRB5*) among the lymphocyte trait variants were significant. In total, 46 (3.6%) unique disease-associated variants and 17 (1.8%) lymphocyte trait variants showed significant division-state interactions (**Supplementary Table 9-10**).

We observed the majority of effects of disease associated variants manifest in early division states compared to late division states. In naive cells, 20 effects (80%) among the disease variants manifested predominantly in early states and 5 in late states. For the lymphocyte trait variants 6 (46%) manifested in early states and 7 in late states. In memory cells, 25 (57%) of the disease variants had a strong effect in early states and 19 in late states, and 4 (40%) of the lymphocyte trait variants did in early states and 6 in late states. For instance, *RPS26,* is especially strongly regulated at early stages of cell division (*RPS26* regulated by the C allele at chr12:56042145; p_interaction_ = 2.0 × 10^−32^; β_early_ = 0.93 and β_late_ = 0.51, **Fig. 2o**), both in naive and memory cells, with a strong reduction of the eQTL effect over time. On the other hand, *HSD3B7* exemplifies a strong late division effect (regulated by the G allele at chr16:30999862 in memory cells; p_interaction_ = 8.5 × 10^−18^; β_early_ = −0.01 and β_late_ = 0.94; **Fig. 2p**).

Together, these analyses demonstrate that the cell division state provides an additional and orthogonal layer of context for resolving genetic regulation of gene expression, complementing conventional activation-time point based analyses. Notably, the proportion of division-dependent eQTLs observed at immune disease–associated loci mirrors that of all eGenes in the dataset (360 genes; 5.6%), suggesting that disease risk variants are not preferentially enriched in proliferation-specific regulatory effects. This observation prompted us to move beyond the analysis of individual gene–variant pairs in division states and ask whether immune disease associations instead converge on other gene expression programmes with distinct division dynamics.

### Gene expression programs highlight selective enrichment of immune disease variants during early division states

Using our division-resolved single-cell dataset, we sought to refine whether disease-associated genetic signals are enriched in discrete pathways that are preferentially active across divisions or cell subpopulations. To capture coordinated transcriptional programs, we applied consensus non-negative matrix factorisation (cNMF) ^48^ to our transcriptomic data. We chose this approach because cNMF can identify positively additive shared and unique gene expression programmes (GEPs), without the requirement of orthogonality (**Fig. 3a**). We identified 23 GEPs that together explained 27% of the total variance in the input dataset that was corrected for the donor and batch effects (**Methods** and **Supplementary Fig. 12**). Programme identities were annotated through integration of transcriptomic, gene set enrichment analysis, and comparison with previously described consensus T cell GEPs (**Fig. 3b, Methods** and **Supplementary Table 11**). GEP usage patterns revealed both strong subpopulation specificity and generic functional programmes shared across cell subpopulations. Five programmes were uniquely enriched in a single subpopulation, including *Treg identity* (Treg), *Cytotoxic granzyme production* (Granzyme⁺/EMRA), *MHC-II antigen presentation* (memory HLA⁺), *Early IFN response* (LA), and an *ER stress* module (Treg cyclin⁺), while sixteen programmes showed usage across multiple subsets in lineage-consistent patterns (**Fig. 3c, Supplementary Fig. 13a**). For example, the *Memory cytokine production* GEP was shared between effector memory and Granzyme⁺/EMRA T cells, while a broadly expressed programme, annotated as *Mitochondria*, was moderately used across all subsets (**Fig. 3c, Supplementary Fig. 13a**).

**Figure 3.**
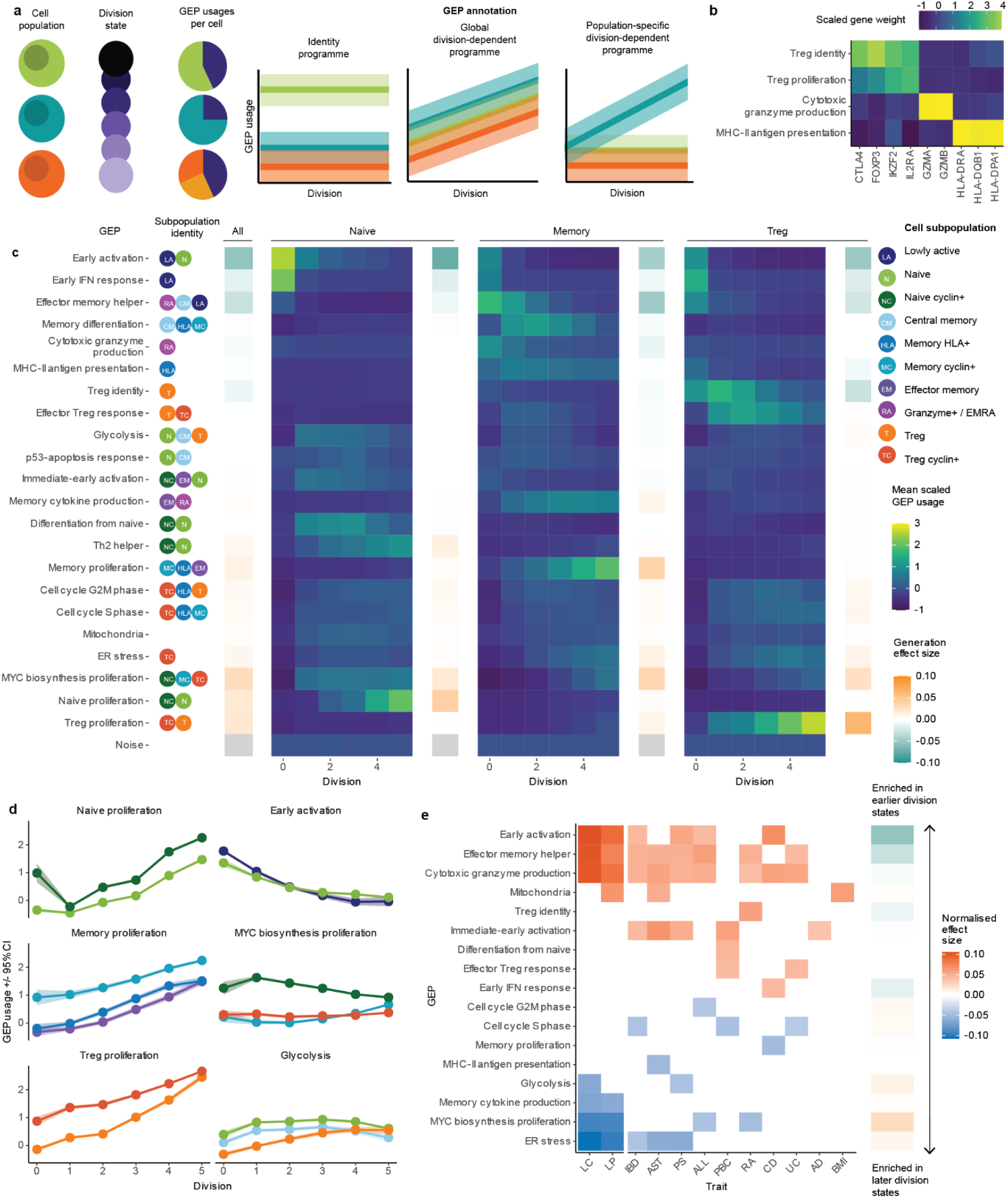
Distinct gene expression programmes underlying T-cell proliferation refine polygenic immune-mediated disease signals. **a.** Schematic of gene expression programmes (GEPs) inferred across proliferative stages in the different cell populations, comprising identity programmes, division-dependent programmes, and population-specific division programmes. **b.** Heatmap showing the highest-weight genes for selected GEPs, including Treg identity, Treg proliferation, cytotoxic granzyme production, and MHC-II antigen presentation. **c.** Heatmap of scaled GEP usage across division states within naive, memory and Treg cells. The circles on the left side indicate cell subpopulations with significant GEP usage, corresponding legend is on the right. The strength and direction of GEP usage associated with the division effect across all cell populations, and within naive, memory and Tregs cells are represented alongside; grey indicates non-significant associations (FDR > 0.05), green and orange represent significant effects, orange - increased GEP usage with divisions, green - decreased GEP usage with divisions. **d.** GEP usage (±95% CI) across cell subpopulations for proliferation-related GEPs, including early activation, MYC-biosynthesis–linked proliferation, and glycolysis. Colours represent cell subpopulations as in the legend for panel c. **e.** MAGMA results for immune-mediated disease and lymphocyte trait SNP enrichment in GEPs. The strength and direction of GEP usage associated with the division states is represented alongside heatmap; grey indicates non-significant association (FDR > 0.05), orange - increased GEP usage with divisions, green - decreased GEP usage with divisions. LC, lymphocyte count; LP, lymphocyte percentage; IBD, Inflammatory Bowel Disease; AST, Asthma; PS, Psoriasis; ALL, Allergic Disease, PBC, Primary Biliary Cholangitis; RA, Rheumatoid Arthritis; CD, Crohn’s Disease; UC, Ulcerative Colitis; AD, Atopic Dermatitis; BMI, body mass index.

The GEPs with subpopulation-specific usage aligned with canonical marker expression. The *Treg identity* and *Treg proliferation* modules were restricted to regulatory T cells and featured high *IL2RA, IKZF2, CTLA4* and *FOXP3* scores (**Fig. 3b**). The *MHC-II antigen presentation* GEP, enriched in HLA⁺ memory cells, was dominated by HLA class II transcripts and CD74, while the *Cytotoxic granzyme production* GEP, enriched in Granzyme^+^/EMRA cells, was dominated by granzyme genes including *GZMA* and *GZMB*. Cell-cycle-associated programmes (*Cell cycle S phase* and *Cell cycle G2M phase*) showed consistent usage across subpopulations and correlated strongly with independent cell-cycle scores (**Methods, Supplementary Fig. 13b**). Three proliferation-associated modules, *naive proliferation*, *memory proliferation* and *Treg proliferation*, exhibited a clear linear increase in usage with division number (p_adj_ < 0.004, |lfc per division| 0.28 - 0.50) and were characterised by high weighting of *MYC*, a central regulator of T-cell growth and division (**Fig. 3c-d; Supplementary Table 11**) ^49^. Notably, several non-proliferation GEPs also exhibited enrichment for division-dependent genes, including *Mitochondria* and GEPs involving naive and memory differentiation and cytokine production, underscoring both the limitations of linear differential expression approaches and the multi-layered, multifactorial nature of transcriptional remodelling across division states. In addition to the annotated proliferation modules, which showed strong positive division effects (p_adj_ < 1×10⁻⁴), division-dependent modelling showed that ∼57% of all GEPs (13/23) exhibited significant changes with division in at least one subpopulation (p_adj_ < 0.05, |lfc per division| > 0.1; **Supplementary Table 11; Supplementary Fig. 14**). Notably, *Early activation, Early IFN response*, and *Effector memory helper* programmes consistently declined across divisions in LA and naive subpopulations, reflecting the expected attenuation of immediate-early transcriptional responses as cells transition from activation into sustained proliferation.

Next, we asked whether polygenic IMD signals preferentially enrich in specific transcriptional programmes. To do so we associated the cNMF spectra scores, which represent the strength of association of a gene to a given program, to GWAS using MAGMA ^50^ (**Methods**). We assessed SNP enrichment across the 23 GEPs for 10 common IMDs (**Supplementary Table 8**), and body mass index (BMI) as a control. We also included lymphocyte count and lymphocyte percentage in the SNP enrichment analysis because they represent quantitative immune phenotypes with a large number of associated genetic variants, expected to show enrichment signal when applied to our proliferation-resolved dataset. Importantly, they are highly relevant to immune-mediated disease pathogenesis, reflecting variation in lymphocyte abundance and activation potential that underpins inflammatory responses.

Across IMDs, SNP enrichment was non-uniformly distributed across defined GEPs, indicating that disease-associated variants preferentially converge on specific functional modules rather than uniformly affecting all expressed CD4⁺ T-cell genes (**Fig. 3e**). At the same time, BMI-associated variants showed no enrichment in immune GEPs and were exclusively associated with the Mitochondria programme, in line with previous reports linking BMI risk variants to mitochondrial dysfunction and the links of this program with metabolism ^51^. Nearly all IMDs showed significant enrichment (p_adj_ < 0.05; effect sizes 0.04–0.10) in *early activation, effector memory helper, and cytotoxic granzyme-production* programmes. These programmes were also preferentially used in early division states (division state β:−0.005 to −0.05; p_adj_ < 0.01), linking disease-associated genetic risk to proliferative effector programmes that engage early following activation (**Fig. 3e**). In addition to these shared patterns, several diseases exhibited programme-specific enrichments consistent with known pathobiology. Rheumatoid arthritis (RA) risk variants were selectively enriched in the *Treg identity* programme, which includes *CTLA4*, the target of the therapeutic antibody abatacept used in RA treatment ^52,53^. In contrast, variants associated with inflammatory bowel disease, asthma, psoriasis, primary biliary cholangitis, and atopic dermatitis were enriched in the *Immediate-early activation* programme characterised by high *IFNG* expression and canonical Th1 regulators ^54^, consistent with the pro-inflammatory cytokine environments observed in these conditions (**Fig. 3c**). Likewise, lymphocyte traits were strongly enriched in activation and differentiation programmes but depleted in modules used in later proliferation stages such as *MYC biosynthesis proliferation* and *ER stress* programmes (**Fig. 3c**). IMD variants were depleted in GEPs with increased usage in late division states such as *ER stress* and GEPs linked to cell cycle, suggesting that while excessive proliferation is a hallmark of inflammation, the genetic risk for IMDs acts predominantly through pathways initiating proliferation and effector differentiation, rather than through the maintenance of cell division itself.

Together, these results show that by resolving cells into discrete division states, this framework distinguishes genetic effects acting on the initiation and licensing of effector responses from those associated with later proliferative maintenance. Such patterns would be obscured in simple time-point comparisons, where asynchronous division masks programme usage and conflates activation with proliferation.

## Discussion

A central challenge in human genetics is to move from association to mechanism by identifying the cellular contexts and biological processes through which disease-associated variants exert their effects. Here we position early CD4⁺ T cell activation as the most likely cellular context causally implicated in complex immune disease, rather than later stages of proliferative expansion. Although proliferation is a defining hallmark of activation, our results suggest that immune disease–associated variants primarily act upstream of proliferation, shaping how CD4⁺ T cells sense, integrate, and interpret stimulatory and co-stimulatory signals that license entry into the cell cycle. Our findings bring new interpretation to studies linking IMD variants to regulation of co-stimulatory pathways ^55,56^, that integrate early stimulation signals to shift activation thresholds and downstream fate decisions. Rather than altering the mechanics of cell division itself, acting at early activation signals, these variants could be altering the likelihood, timing, or stability of entry into activation states that are competent for cell-cycle entry and influencing the trajectory of effector differentiation.

Our analyses reveal that gene expression is context dependent throughout the proliferative trajectory and genetic effects on gene regulation can manifest at specific stages of proliferation. As regulatory effects become increasingly pervasive ^57,58^, identifying the cellular contexts in which these effects manifest becomes critical for anchoring genetic associations in biological mechanisms. A striking example of this is *RPS26,* a pleiotropic gene associated with numerous immune diseases and cellular traits ^7,9,59–65^, a hotspot for *trans*-eQTL ^63,65^, co-expression-QTL ^61,62^, and Mendelian randomization hits for 81 traits ^64^. We observed a strong genetic effect manifesting especially at early division states, suggesting that the mechanism underlying these findings may well be driven by early activation and the entry into the cell cycle. *RPS26* knockout mice have reduced numbers of peripheral CD4^+^ and CD8^+^ T cells as well as a smaller thymus, an increase in apoptosis, but the remaining T cells do not have impaired proliferative capacity ^59^. These results support the link to early activation and cell cycle entry, likely not to be only specific to T cells but more broad across several cell types given its ubiquitous expression and high pleiotropy ^64,66^. Our results shed new light on the biological interpretation of this gene across different cellular and stimulatory contexts. For example, *RPS26* was identified as a cell type specific response eQTL with especially strong effect in B cells from PBMCs that were transcriptionally profiled 24h after exposure to different stimuli ^7^. B cells can proliferate rapidly after antigenic stimulation, rating about 1 division every 6 hours ^67^, therefore, the detection of B cell effect could be attributed to this cell type being faster to proliferate, manifesting in detection of the *RPS26* effect which is regulated at early division states. Together, these observations indicate that proliferation is not merely a downstream consequence of activation, but a functional axis along which regulatory variation manifests.

There is a growing interest in understanding context-dependent dynamic eQTLs by modelling genetic effects at the level of individual cells, which can capture fine-grained cellular heterogeneity ^8,9,26–28^. In particular, interaction models that incorporate genotype and a quantitative cellular state enable explicit testing of context-dependent regulation. We observed systematic inflation of genotype–cell-state interaction statistics when using the previously proposed NBME model. While main genotype effects were well-calibrated, interaction terms were not. Incorporating a donor-level random slope substantially improved model calibration and attenuated association statistics in the permuted data, suggesting that the original model is prone to spurious interaction signals. Although evaluated here only for division state, these findings underscore the need for careful calibration when applying single-cell–level eQTL interaction analyses in other settings.

In this work we demonstrate that high-level cellular phenotypes, such as division state, can be inferred directly from transcriptomic profiles and used to stratify genetic analyses at scale. By training *CellDivider* to predict division state from single-cell transcriptomes, we show that proliferative heterogeneity can be quantified across individuals in independent datasets, revealing substantial inter-individual variability that is more pronounced in memory than naive CD4⁺ T cells. Importantly, as complex trait polygenic signals converge on specific gene programmes and pathways ^68,69^, it underscores the need to map genetic effects onto cellular phenotypes rather than isolated molecular readouts. For example, high-content image-based cell phenotyping studies in metabolic disease have demonstrated how profiling high-dimensional cellular phenotypes and correlating them with genetic variation can link risk variants to disease-relevant cellular phenotypes ^70^. Therefore, scalable approaches that enable inference of cellular traits at population scale are critical for identifying how complex trait variants perturb cellular behaviour and for defining the genetic regulators of cell phenotypes. While the genetic dataset used here is of modest size, applied to larger cohorts, this framework offers a path towards propagating genetic signal from variant to gene, from gene to programme, and ultimately to cellular phenotype.

Our analyses show that the transcriptional consequences of proliferation differ substantially across CD4⁺ T cell subsets. Naive T cells undergo the most extensive division-associated transcriptional remodelling, coupling proliferation to broad activation, metabolic, and signalling programmes, whereas memory and regulatory T cells exhibit attenuated and more selective responses. This pattern is consistent with a model in which naive cells require extensive reprogramming upon first activation, while antigen-experienced cells, already primed by prior activation, constrain transcriptional plasticity during proliferation ^71,72^. This division and cell type specific gene expression regulation is also influenced by the genetic variation. An example is *RFLNB*, a gene encoding an actin-binding protein. Although there are no prior reports directly implicating differential regulation of *RFLNB* between naive and memory T cells, substantial differences in actin organisation, cytoskeletal dynamics, and cellular mechanics are well established as key determinants of their distinct migratory and activation behaviours ^73^. Our findings suggest that genetic regulation of *RFLNB* may contribute to these differences, particularly at later stages of division in memory cells, when proliferative responses are decoupled from early activation programmes. The presence of subset-specific division effects further indicates that cell identity conditions how proliferative programmes are deployed and are subject to genetic regulation.

We also recognise several limitations of our study. First, the genetic analyses were performed in a cohort of 119 individuals, which limits power to detect variant effects on cellular phenotypes and interaction effects. The lack of strong genetic drivers of *in vitro* proliferation suggests that variation in cell division is driven by a highly polygenic architecture with many small-effect variants, in line with previous work ^74^, and influenced by environmental factors. We expect that larger cohorts will reveal a greater number of genetic signals. Although we validated *CellDivider* in independent datasets generated using similar activation systems, differences in stimulation strength, and experimental parameters remain, and should be considered when applying *CellDivider* to *in vivo* or PBMC-derived datasets.

Second, given the limited power and high computational burden of single cell level mixed linear models, we only opted to test top *cis*-eQTL main effect variants and GWAS top variants for division state interaction. It is likely that we capture only a subset of proliferation-linked regulation, and we therefore expect division-specific genetic effects to be more abundant than estimated here. A clear example is *TAPBPL*, whose genetic regulation is not apparent when expression is analysed across pooled activated cells. However, resolving proliferation reveals a division-dependent regulatory effect in naive CD4⁺ T cells, which is most pronounced at later division states. This pattern is biologically consistent with a role for *TAPBPL* in constraining T cell expansion after initial activation ^75^ and illustrates how context-resolved analyses can uncover regulation of key immune inhibitory pathways that is otherwise masked. Future work could reveal more such effects in a genome wide scan of division state eQTLs.

Finally, our study examines proliferation in an *in vitro* setting using antigen-invariant T cell stimulation. Under these conditions, we assume that cells have an equal probability of activation over the course of the assay, resulting in a naturally unsynchronised population spanning multiple division states. This design enables us to capture division-dependent transcriptional changes, but it also relies on the assumption that activation timing is largely stochastic rather than driven by stable intrinsic differences between cells. Moreover, the *in vivo* environment, including antigen specificity, cytokine milieu, and metabolic constraints, is likely to shape proliferative and differentiation trajectories in ways not captured here. Longer-term differentiation processes may also extend beyond the five-day window examined in this study.

In summary, we establish cell division as a regulatory axis in CD4⁺ T cell biology, showing how modelling dynamic cellular contexts can sharpen variant-to-function inference. By providing a robust, extensible baseline for interpreting genetic effects, it lays the groundwork for broader application to other immune cell types or models that track division history in quiescent and tissue-resident cells. This approach enhances interpretation of single-cell transcriptomic data, deepens understanding of immune regulation and immune-mediated disease pathobiology, and supports context-aware genetic studies that help prioritise disease-relevant pathways and cell states for therapeutic design.

## Methods

### Donor samples

Peripheral blood mononuclear cells (PBMCs) were obtained from four healthy adult females (mean age 38.5 years, s.d. 3.4) via leukopaks supplied by BioIVT. Samples were collected under informed consent and used in accordance with IRB/EC-approved protocol 15/NW/0282. PBMCs were isolated using Ficoll-Paque PLUS density gradient centrifugation and cryopreserved within 24 hours of collection.

### Experimental design

Cryopreserved PBMCs were thawed, rested overnight in complete RPMI (10% FBS, 1% PenStrep, 1% L-glutamine), and CD4⁺ naive, memory, or Treg cells were isolated using EasySep™ kits (EasySep™ Human Naive CD4⁺ T-cell Isolation Kit, #19555, EasySep™ Human Memory CD4⁺ T-cell Enrichment Kit, #19157, EasySep™ Human CD4⁺CD127_low_CD25⁺ Regulatory T-cell Isolation Kit, #18063). Cells were profiled across three batches, with each batch comprised one donor representing one CD4⁺ T-cell population. One population was represented by more than one donor across batches to minimise confounding effects.

### CellTrace Violet staining and stimulation

Cells were stained with CellTrace Violet (CTV, ThermoFisher) at 1 µL /2 mL, incubated at 37 °C for 20 minutes with gentle agitation every 5 minutes, quenched in cold medium, washed, and resuspended in StemPro medium. Isolated Tregs were supplemented with recombinant IL-2 (50 ng/mL). All subsets were stimulated with ImmunoCult™ Human CD3/CD28/CD2 Activator (12 µL per million cells) and plated at 1M cells/mL. Media was replenished on day three and cells harvested on day four.

### Cell sorting

Cells were stained with 7AAD for live/dead discrimination and filtered through 30 µm mesh prior to sorting on an Invitrogen BigFoot sorter. Six division states were identified based on CTV intensity, and ∼300,000 cells were sorted per peak and per donor into RPMI supplemented with 30% FBS and kept on ice until CITEseq (TotalSeq™-C Human Universal Cocktail; BioLegend, 399905) and cell hash staining.

### CITE-seq staining

Sorted cells were incubated with the TotalSeq-C antibody cocktail (BioLegend, 399905) and TotalSeq-C cell hashing antibodies (Biolegend) following the manufacturer’s instructions. Each vial contained 500,000 cells, generated by pooling cells from all donors in equal proportions across distinct division states. To minimise non-specific binding, cells were first incubated with Fc block before staining. Cells were then incubated with the CITE-seq cocktail and corresponding antibodies at 4 °C for 30 minutes, followed by three washes with cell staining buffer prior to 10x library preparation.

### Validation dataset

An independent single-cell validation dataset comprising 25,694 activated CD4⁺ T cells sorted by CTV-labelled division number was generated from PBMCs from one of the donors previously cryopreserved. Cryopreserved PBMCs were thawed and rested overnight in complete RPMI (cRPMI: RPMI 1640 (ThermoFisher, 52400041) supplemented with 10% FBS (Sigma-Aldrich, F9665), 1% PenStrep (ThermoFisher, 15140122) and 1% L-glutamine (Sigma-Aldrich, G7513)). Memory CD4⁺ T cells were isolated using the EasySep™ Human Memory CD4⁺ T-cell Enrichment Kit (StemCell Technologies, 19157). CTV staining was performed as described for the main dataset prior to stimulation with Dynabeads Human T-cell Activator (anti-CD3, anti-CD28; ThermoFisher, 11131D) at a 2:1 cell:bead ratio. Cells were counted and resuspended in cStemPro supplemented with IL-2 (455 units/mL IL-2, Biotechne, 10453-IL-010) at 1M cells per ml.

Sorting was performed as described for the main dataset at 16 hours, day 3 and day 5 post-activation, alongside a resting sample collected at 16 hours. For CITE-seq staining, the TotalSeq-C lyophilised antibody cocktail (BioLegend, 399905) was equilibrated at 21 °C for 5 minutes and centrifuged at 10,000 g for 5 minutes. Vial contents were rehydrated with 27.5 µL cell staining buffer, vortexed and incubated for 5 minutes, then transferred to LoBind protein Eppendorf tubes (ThermoFisher, 90410) and centrifuged at 14,000 g for 10 minutes at 4 °C. The supernatant was diluted 1:6 in cell staining buffer, vortexed and mixed thoroughly. Fc blocker (ThermoFisher, 422302) was added at 5 µL per million cells and incubated at 4 °C for 10 minutes. A total of 750,000 cells were stained in a 1:12 dilution of the CITE-seq cocktail and incubated at 4 °C for 30 minutes, followed by three washes in cell staining buffer. Cell viability was monitored using AO/PI staining assessed by Cellometer Spectrum (Nexcelom).

Cell hashing antibodies were used to assign division states and time points. For the 16-hour sample, cell hashing antibody (1 µL per ∼750,000 cells) was added directly to the CITE-seq cocktail. For the day 3 time point, cell hashing was performed as an independent staining step (including three washes) prior to CITE-seq staining. For the day 5 time point, CITE-seq staining was performed before sorting, and cell hashing was carried out on the sorted cells in a separate staining step.

All subsequent steps for 10x Genomics library preparation, sequencing and downstream processing were performed identically to the main experimental dataset.

### GEM generation, library preparation and sequencing

Single-cell suspensions were loaded at ∼25,000 cells per 10x Genomics Chromium reaction. Libraries were generated using Chromium Next GEM Single Cell 5’ v2 chemistry following the manufacturer’s protocol ^76^, 10X Genomics). Both gene expression (GEX) and antibody-derived tag (ADT) libraries were constructed and sequenced on Illumina NovaSeq S4 lanes (150 bp paired-end). Sequencing depths targeted >50,000 reads per cell for GEX and >10,000 reads per cell for ADT, using a 5:1 GEX:ADT ratio.

### Data processing

Sequencing base calls were demultiplexed into FASTQ files with Cell Ranger (v6.1) ^76^. Reads were aligned to GRCh38 (GENCODE v32) using Cell Ranger multi, which generated single-cell feature matrices and cell hashtag sample demultiplexing. Donor deconvolution was performed using natural genetic variation with Vireo and Souporcell, with consensus donor assignments verified via PLINK ^77–79^

Hashtag demultiplexing was refined in Seurat (HTOdemux) using centred log-ratio (CLR) normalisation and the clara k-medoids clustering algorithm ^80^. The positive classification threshold for each hashtag was set at the 0.99 quantile of the inferred negative distribution. Multiplets and unassigned droplets were excluded from downstream analyses.

### Quality control and filtering

Cells were filtered using batch-specific thresholds to remove low-quality or non-CD4⁺ populations. Low-quality cells were defined as those with low read depth or gene counts (≤ 2–3 median absolute deviations (MADs) from division 0 benchmarks), high mitochondrial read fraction (> 15% or > 3 MADs), or outlier ADT counts (> 3 MADs). Non-CD4⁺ clusters were identified via principal component analysis (PCA), louvain clustering and UMAP, and excluded based on transcriptomic and surface protein markers: dendritic cells (CD86⁺), B cells (CD19⁺), monocytes (CD36⁺), CD8⁺ T cells (CD8A⁺), γδ T cells (TCRγδ⁺), and NK cells (CD16⁺).

Across all samples, nearly half of detected (n=73,305) UMI’s were removed; 41,859 (57%) were removed due to low read depth or gene count, 25,705 cells (35%) for high mitochondrial content, 750 cells (1%) for outlier ADT counts, and 4,991 cells (7%) as non-CD4⁺ contaminants.

After cell-type annotation (see below), cells were filtered to ensure concordance between the isolation strategy and post hoc cell-type annotation, thereby maintaining consistent culture conditions across populations. Specifically, transcriptomically annotated naive cells (naive and naive cyclin⁺, *TCF7+PLAC8+CCR7+*) were required to have been initially isolated as CD4⁺CD45RA⁺CD45RO⁻ naive T cells (EasySep™ Human Naive CD4⁺ T-cell Isolation Kit, 19555). Memory cells (central memory, memory HLA⁺, memory cyclin⁺, effector memory, and granzyme⁺/EMRA) were required to have been isolated as CD4⁺CD45RA⁻CD45RO⁺ cells (EasySep™ Human Memory CD4⁺ T-cell Enrichment Kit, 19157). Treg cells (Treg and Treg cyclin⁺) were required to have been isolated as CD4⁺CD127_low_CD25⁺ cells (EasySep™ Human CD4⁺CD127_low_CD25⁺ Regulatory T-cell Isolation Kit, 18063) and to express *FOXP3* and *IL2RA* at the transcriptomic level. Cells with discordant isolation and annotation labels were excluded. This yielded 27,151 naive, 23,269 memory, and 16,339 Treg single cells used for downstream analysis.

### Normalization, clustering and cell type annotation

Seurat version 5 was used for single-cell processing ^81,82^. Cell cycle phase was inferred using Seurat’s CellCycleScoring function and the cc.genes.updated.2019 gene set ^83^, generating S and G2/M scores for each cell, phase classification (G1, S, or G2/M) was assigned based on the relative weighting of these scores.

Remaining cells were re-normalised using SCTransform log-normalise (Seurat v5), and scaled to regress out cell cycle and mitochondrial effects. 2,000 highly variable features were identified per batch, which was reduced to 1922 following exclusion of TCR variable region transcripts.

Batch, donor, and cell cycle effects were integrated using Harmony ^84^ to ensure robust clustering across datasets. Dimensionality reduction was performed using principal component analysis (PCA) followed by shared nearest neighbour (SNN) graph construction (k = 20, 30 PCs). Clustering was conducted with the Louvain algorithm at resolutions between 0.2 and 0.5, with 0.4 deemed to be optimal. The ten clusters were manually annotated using canonical transcriptomic markers with supporting surface protein profiles ^9^, resolving populations into LA (*KLF2, IL7R, TCF7*), naive (*TCF7, PLAC8, CCR7*), naive cyclin⁺ (*TCF7, PLAC8, CCR7, CCNB1, CDC20*), central memory (*CCR7, MPRS26, S100A6, ITGB7, VIM*), memory HLA⁺ (*MPRS26, ITGB7, HLA-DRA, HLA-DPA1, HLA-DPB1*), memory cyclin⁺ (*MPRS26, ITGB7, CCNB1, CCNB2, CDC20, CDKN3*), effector memory (*S100A6, ITGB7, VIM, CCL5, CCL4, CCL3*), Granzyme^+^/EMRA (*ITGB7, VIM, CCL5, CCL4, CCL3, GNLY, GZMB, GZMA, NKG7*), Treg (*FOXP3, IL2RA, CTLA4, IKZF2*) and Treg cyclin⁺ (*FOXP3, IL2RA, CTLA4, IKZF2, CCNB1, CCNB2, CDC20, CDKN3*) subsets (**Supplementary Fig. 2a-c**).

A total of 7%, 10% and 48% cells isolated by naive, memory and Treg EasySep kits were annotated to contrast to their isolation kit respectively (ie. Treg cells isolated and cultured as naive cells), reflecting known limitations of antibody-based T-cell population isolation (**Supplementary Fig. 15**).

### Variance partitioning

To quantify the contribution of experimental and biological factors to gene expression variability, variance decomposition was performed using the *variancePartition* package ^85^. In brief, *variancePartion* models the normalized gene expression of each gene as a mixed linear model, where categorical variables are treated as random effects. For single cell analysis we used the *Seurat* SCTransform data slot, for pseudobulk *Seurat* NormalizeData was used which produces log(cp10k+1). All quality-controlled cells NMT were included, and variance was modelled across batch, donor, cell population, and division state (treated as a categorical variable) for the 2,000 most variable genes. The linear mixed model was specified as:

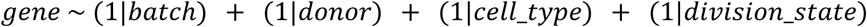

For the pseudobulk analysis the same model was applied.

To correct for cell-cycle effects, we applied the same model, but used the *Seurat* SCTransform specifying the G2M and S scores as covariates and supplied the scaled residuals to *variancePartition*. For the pseudobulk we calculated the G2M and S scores on the pseudobulk in the same manner as described above. This should represent a conservative way to correct for the cell-cycle effect, especially given division and cell-cycle are strongly correlated processes.

### Differential expression analysis

Pseudobulk expression profiles were generated per donor, population, and division state. Genes expressed in fewer than 25% of single cells were excluded. In total 7,833 genes were included for differential expression analysis. Differential expression analysis was performed using *DESeq2* ^86^, controlling for donor and mean cell cycle scores (S and G2M), by taking an average of the cell cycle scores per pseudobulk group. The naive population served as the baseline level. Interaction terms were included to test for division–subpopulation effects, according to the model:

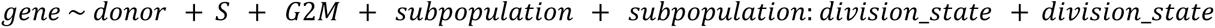

Log₂ fold changes were shrunk using *ashr* ^87^. Genes or proteins were considered significantly differentially expressed at p_adj_ < 0.05 and |β| > 0.1 (equivalent to |lfc| > 0.1 per division state) for the interaction coefficients (*subpopulation:division_state* and *division_state*).

To identify divergent genes exhibiting significantly different division-dependent dynamics across subpopulations, the subpopulation interaction term was extracted. Divergent genes were defined as those with p_adj_ < 0.05 and |β| > 0.1.

### Gene set enrichment analysis

Gene lists generated from differential expression analysis were used for gene set enrichment analyses. Analysis was implemented in sc-blipper ^88^, a Nextflow pipeline for post-processing scRNAseq datasets and performing gene set enrichment tests using GSEA (log2 fold changes for all genes) and ORA (significant genes only) through R packages fgsea and fora ^89^. All 7,833 genes tested in differential expression as the background set for enrichments. As reference datasets we used gmt files from various sources, but primarily from msigdb ^90^ (**Supplementary Table 12**). Gene sets with fewer than 5 genes and more than 2000 were omitted. Exact files used are available at https://github.com/TrynkaLab/sc-blipper/tree/main/assets/gene_sets/symbols. The false discovery rate for enrichments was determined by Benjamini-Hochberg correction ^91^ over all tested reference datasets, within each condition and enrichments with FDR < 0.05 were considered significant. A matrix of regression values was used to identify the division-dependent gene processes per cell state. Significance was evaluated by Benjamini-Hochberg adjusted p-values per test over all tested databases as < 0.05.

### Development of the *CellDivider* predictive model

We benchmark three methods (elastic net regression, eXtreme Gradient Boosting ^92^ [XGBoost] and a multi-layer perceptron [MLP]) and evaluated five different input features (normalised gene expression of 2000 HVGs, top 30 Harmony PCs, 23 cNMF GEP usages, 30 scVI latent variables and scVI-adjusted gene expression of 2000 HVGs).

### Input features

*Normalised gene expression*: to prepare normalised gene expression, we applied SCTransform ^93^ to the single-cell datasets by regressing out batch, donor, percentage mitochondrial reads and percentage ribosomal reads.

*Harmony PCs*: we integrated the main and validation datasets by applying the RunHarmony function in the harmony R package ^84^ to the top 30 PCs based on the normalised gene expressions above.

*cNMF GEP usages*: the details of cNMF GEP derivation are described in the ***cNMF*** section later.

*scVI latent variables and adjusted gene expression*: we trained a scVI model of one layer and 30 latent variables ^94^ while adjusting for donor, batch, percentage mitochondrial reads and percentage ribosomal reads. Latent variables were obtained by the ‘get_latent_representation()’ method. Adjusted gene expression was obtained by the ‘get_normalized_expression()’ method and log-transformed.

### Machine learning models

We formulated division-state prediction as a regression problem, treating division state as a continuous (real-valued) rather than a discrete (integer-valued) value to more accurately capture the inherent uncertainty in the predictions, with the aim of maximising power in the following regression analyses (eQTL analyses). The elastic net regression models were implemented using the scikit-learn python package ^95^. The XGBoost models were implemented using the xgboost python package ^96^. The MLP models were implemented using the pytorch python package ^97^ as a 3-layer MLP with a ELU activation ^98^ and with a final Softplus layer to enforce non-negative outputs. The weights were optimized by nn.L1loss() (mean absolute error) using ADAM ^99^, since we formulated the task as a regression problem

### Model benchmarking and optimisation

For model benchmarking and optimisation, we used the main dataset (86,760 cells) for hyperparameter tuning and an independent test dataset (25,694 cells) to evaluate generalisability. We optimised hyperparameters of the models using 5-fold cross-validation with a stratified train-validation split on the main dataset with Optuna ^100^ The hyperparameters were optimised with the aim of maximising the Pearson correlation coefficient between the observed and predicted division states. The hyperparameters selected for the final MLP model are listed in **Supplementary Table 13**. The final model, using these optimised hyperparameters, was trained on the full main dataset. The generalisability of the final model was assessed on the completely held-out test dataset, where samples were held-out during training.

During model training for the population-scale dataset (655,349 cells from 119 individuals) ^9^, given the increased number of cells and potential complexity in the combined dataset, we used a one-layer and 100 latent variable scVI model to prepare the input features. Given that the population-scale dataset consisted of naive and memory cells, we restricted the training set to these cell types.

To assess which input features (scVI-adjusted gene expression) contribute most to the prediction, we applied the integrated gradients method ^101^ to the independent test dataset, yielding a cell-by-feature attribution matrix. The baseline for the attribution computation was set to the mean of the undivided cells (division state = 0). The attribution per gene was computed by taking the mean of the attributions across all cells.

### Deriving proliferation statistics

To determine the predicted proliferation statistics for each donor in the population-level dataset, we rounded the predicted division state to the nearest integer, and for each donor, counted the number of cells in each division. Using these numbers we then calculated the following statistics for each donor which were defined as follows:

Proliferation index:

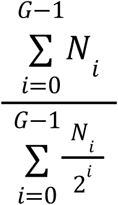

Where *G* is the total number of division states observed, i represents the division state and *N_i_* is the number of cells observed in division state *i*. It can be viewed as the average number of cells that a founding cell became

Division index:

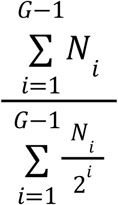

Where *G* is the total number of division states observed, *i* represents the division state and *N_i_* is the number of cells observed in division state *i*. It can be viewed as the average number of cells that a dividing cell became, as this does not account for the undivided state.

Percent divided:

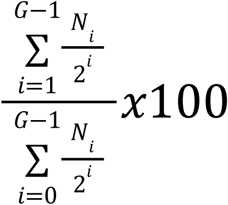

Where *G* is the total number of division states observed, *i* represents the division state and *N_i_* is the number of cells observed in division state *i*. It represents what percentage of the founding population underwent division. This has been implemented in the function ‘get.prolif.stats()’ and is available on github: https://github.com/TrynkaLab/ProliferationAnalysis.

### GWAS of predicted proliferation statistics

QC of the genotypes used in the predicted proliferation GWAS and eQTL mapping has been previously described in Soskic et al. ^9^. We mapped GWAS effects for 40h and 5 day timepoints for proliferation index, division index and percentage divided in both naive and memory population. GWAS effects were modelled using TensorQTL in *trans* mode, supplying the single cell processing batch as a covariates. Variant effects are reported with b38 positions and effect alleles represent the allele with dosage 2.

### Single-cell division-dependent eQTL analysis

To test division-dependent changes in genetic regulatory effects, we modelled single-cell-level gene expression using NBME models.

### Previously proposed NBME model

We initially adopted the previously proposed NBME model composed of fixed effects and random intercepts for donor and batch ^27^. Specifically, to test the genotype main effect, we used a NBME model:

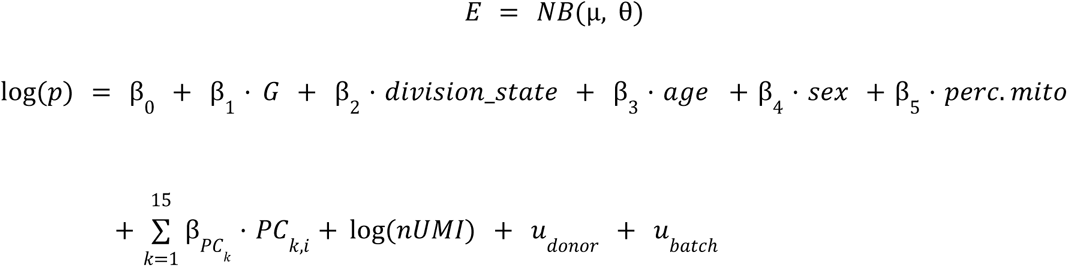

where:

*E*: the read counts of the gene for the cell,
θ: the dispersion parameter of the negative-binomial distribution,
*G*: the genotype dosage of the donor,
*division*_*state* : the continuous division state value,
*age*: the age of the donor,
*sex*: the sex of the donor,
*perc*. *mito*: the mitochondrial transcript percentage for the cell,
*PC*_*k*_: the *k*th expression principal component,
log(*nUMI*): the offset term accounting for library size differences,
*u* _*donor*_, *u* _*batch*_ : the random intercepts for donor and batch, respectively,

We fit the model using the ‘glmer.nb’ function in the lme4 R package with options nAGQ = 0 and ‘nloptwrap’ optimiser. To assess the significance of the genotype effect term β_1_ · *G*, we compared the full model to the reduced model without the genotype effect term using the likelihood ratio test.

When testing the interaction between genotype and division state, we added the interaction term β · *division*_*state*) to the model:

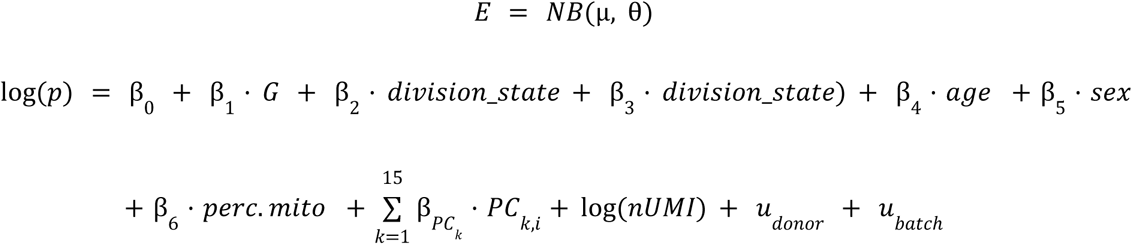

We fit the model in the same procedure and assessed the significance of the interaction by comparing the full model to the reduced model without the interaction term using the likelihood ratio test.

### Extended NBME model

To account for the between-donor heterogeneity in the division state effect size, we extended the NBME model by adding a donor-level random slope:

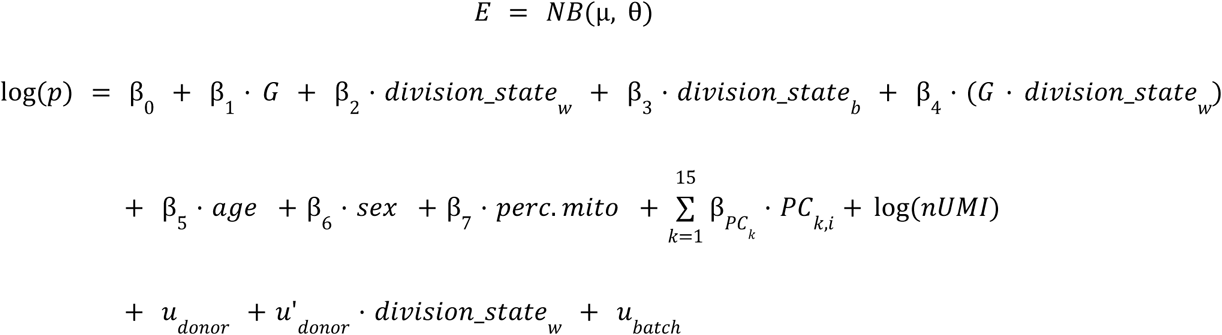

where:

*division*_*state*_*w*_: the within-donor (cell-level) component of the division states,
*division*_*state*_*b*_ : the between-donor component of the division states (donor-level mean),
*u*’_*donor*_: the random slope regarding *division*_*state*_*w*_ for donor.

In this model, to disentangle cell-level from donor-level associations, the division_state score was decomposed into within-donor (*division*_*state*_*w*_) and between-donor (*division*_*state*_*b*_) components using a standard within-between decomposition ^102^. We fit the model using the ‘glmer.nb’ function in the lme4 R package with options nAGQ = 0 and ‘nloptwrap’ optimiser. To assess the significance of the interaction term β_4_· (*G* · *division*_*state*_*w*_), we compared the full model to the reduced model without the interaction term using the likelihood ratio test. To filter out potential model misspecification, we removed tests with an abnormally low (θ<0.1) estimated dispersion parameter. We applied Storey’s *q*-value method ^103^ for multiple testing correction.

For the significant variant–gene pairs, we computed the representative genotype effect size in the early division states 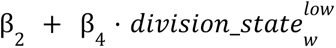 and that in the late division states 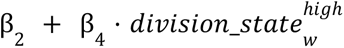, where 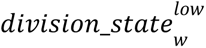 and 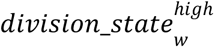 are 2.5 percentile and 97.5 percentile of *division*_*state*_*w*_, respectively.

### cNMF

To define the gene expression programmes (GEPs) underlying our T-cell proliferation dataset in a data-driven way, we applied consensus non-negative matrix factorisation (cNMF) using the Python package cNMF v1.6.0 ^48^. The input read count matrix of the top 2,000 highly variable genes was adjusted for donor and batch effects using the preprocess_for_cnmf function in the package. The number of derived GEPs (*K*) is a hyperparameter specified before running the algorithm. We ran cNMF with 100 iterations for each *K* = [3, 4, 5, 6, 7, 8, 9, 10, 11, 12, 13, 14, 15, 17, 19, 21, 23, 25, 27, 29, 30, 35, 40, 45, 50, 55, 60, 65, 70, 75, 80, 85, 90, 95, 100]. Based on the trade-off between the two benchmarking statistics, stability and proportion of variance explained, we selected *K* = 23 (**Supplementary Fig. 12**).

### Annotation of cNMF programs

The 23 cNMF-derived GEPs were annotated using a combination of subpopulation-specific usage testing, functional enrichment, marker-based inspection, and quantitative association analyses (**Supplementary Table 11**).

To identify cell subpopulations exhibiting preferential usage of individual GEPs, mean GEP usage per donor and subpopulation was calculated. A one-way ANOVA was performed across subpopulations for each GEP, and those with significant differences following Benjamini–Hochberg correction (FDR < 0.05) were further assessed using paired one-vs-all t-tests to determine whether a specific subpopulation displayed significantly higher usage relative to the other subpopulations. Significant differences were validated by visual inspection of subpopulation-stratified usage distributions.

Functional annotation was performed using sc-blipper ^104^ applying both over-representation analysis (ORA) and gene set enrichment analysis (GSEA) as described above under **Gene set enrichment analysis**. For input into GSEA the cNMF spectra scores were used, which represent the relative strength of the contribution of the gene to the program. For ORA the top 50, 100, 250 and top 500 genes for each program were determined by ranking the spectra scores. The set of 7,833 genes tested in differential expression was used as the background set for enrichments. Metabolic programmes were further annotated using curated metabolic pathway gene lists from scCellFie ^105^ and the gene usages.

To link GEPs to division-state and cell-cycle dynamics, Pearson correlation coefficients were computed between GEP usage and (i) division number and (ii) Seurat-derived S- and G2M-phase scores. GEPs showing strong positive or negative linear associations (|r| above the analysis-specific threshold, supported by visual inspection) were annotated as division-linked or cell-cycle-linked programmes.

Initial manual labels were then refined using the highest-weighted genes and proteins contributing to each programme, prioritising transcription factors, cytokines, and well-characterised markers of T-cell activation and lineage state.

Finally, to contextualise the programmes within established NMF-derived signatures, we compared our GEPs to the curated reference programmes reported by ^48^ using the Jaccard similarity index, enabling alignment with previously described canonical gene programmes.

To assess GEP–division relationships, we modelled division-dependent changes in GEP usage by computing the average scaled usage (scaled per GEP) per donor and division state, and fitting the linear model:

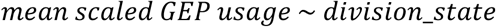

These analyses were performed at three levels: (i) all cells combined, (ii) per subpopulation, and (iii) restricted to the subpopulation–GEP pairs showing significant preferential usage from the ANOVA/t-test framework. For narrative interpretation, division-dependent trends were reported only for the subpopulation–GEP pairs with significant subpopulation-specific usage, ensuring that division-linked effects were assessed in biologically relevant contexts.

### Genetic enrichment of cNMF programs

To associate GWAS summary statistics to cNMF programs we applied MAGMA v1.10 ^50^. In brief, MAGMA converts variant level genome wide summary statistics into a gene level score using variants 50kb upstream and 10kb downstream around the gene body. It then calculates gene-gene relationships for these scores likely to come from LD using an LD reference panel. We used 1000G European individuals ^106^ and variants filtered at MAF<0.05. We calculated gene scores for 19,790 protein coding genes (excluding genes in the extended HLA region chr6:25726063-33400644 (b38) ^107^) in Ensembl version 114 and 33 GWAS traits listed in **Supplementary Table 8**. As responses we used the 23 cNMF gene spectra scores as input for MAGMA -gene-covar which runs associations as separate regression models ^50^. Regressions were run for 7,833 genes identified as expressed. This workflow has been implemented in the sc-blipper Nextflow pipeline described above.

## Code and data availability

The analysis scripts supporting this paper are available at: https://github.com/TrynkaLab/CellDivider and the code for the prediction model & evaluation is available at: https://github.com/TrynkaLab/celldivider_mlp. The code for running gene set enrichment can be found in: https://github.com/TrynkaLab/sc-blipper

Raw sequencing data supporting this study has been deposited in the European Genome-phenome Archive (EGA) and will be made available upon publication. Processed single cell Seurat objects and model weights have been deposited in EBI BioStudies under accession S-BSST2659 (DOI: <u>10.6019/S-BSST2659</u>).

The *Soskic et al.,* dataset used for eQTL mapping in this study has been previously deposited in EGA with accession number EGAD00001008197. Genotypes have been deposited in the European Genome-phenome Archive with accession number EGAD00010002291 and processed single-cell data are available at https://trynkalab.sanger.ac.uk.

## Acknowledgements

We thank members of the Trynka laboratory for helpful discussion and feedback throughout the project. In particular, we thank Florence Lichou, Reem Satti, Marta Perez-Alcantara, Luca Stefanucci, Gareth Griffiths, Anna Lorenc and Thomas Vanderstichele for providing support and discussing results.

We would like to express our thanks to Bee Ling, Jade Joseph and Jennie Graham for her assistance with running and maintaining the Flow Cytometers used in this study. We would further like to thank the DNA pipelines and Human Genetics Informatics teams at the Wellcome Sanger institute for their continued assistance in sequencing and running and maintaining the high performance compute infrastructure.

## Author contributions

M.E.G. was involved in: Investigation, Formal analysis, Methodology, Visualization, Writing - original draft; K.S. was involved in: Investigation, Conceptualization, Formal analysis, Methodology, Visualization, Writing - original draft; K.L. was involved in: Formal analysis, Methodology, Writing – review & editing; T.R. was involved in: Methodology; Z.K. was involved in: Formal analysis, Methodology; X.I.S. was involved in: Formal analysis, Methodology; M.L. was involved in: Methodology, Supervision; B.S. was involved in: Conceptualization, Writing – review & editing; C.P.J. was involved in: Conceptualization, Resources, Project administration, Supervision; O.B.B. was involved in: Conceptualization, Formal analysis, Data curation, Methodology, Software, Supervision, Visualization, Writing - original draft; G.T. was involved in: Conceptualization, Funding acquisition, Project administration, Resources, Supervision, Visualization, Writing - original draft. All roles defined according to the Credit taxonomy: https://credit.niso.org/

