## Supplementary Figures for "Division state reveals hidden genetic regulation during T cell activation and identifies immune disease–linked gene programmes"

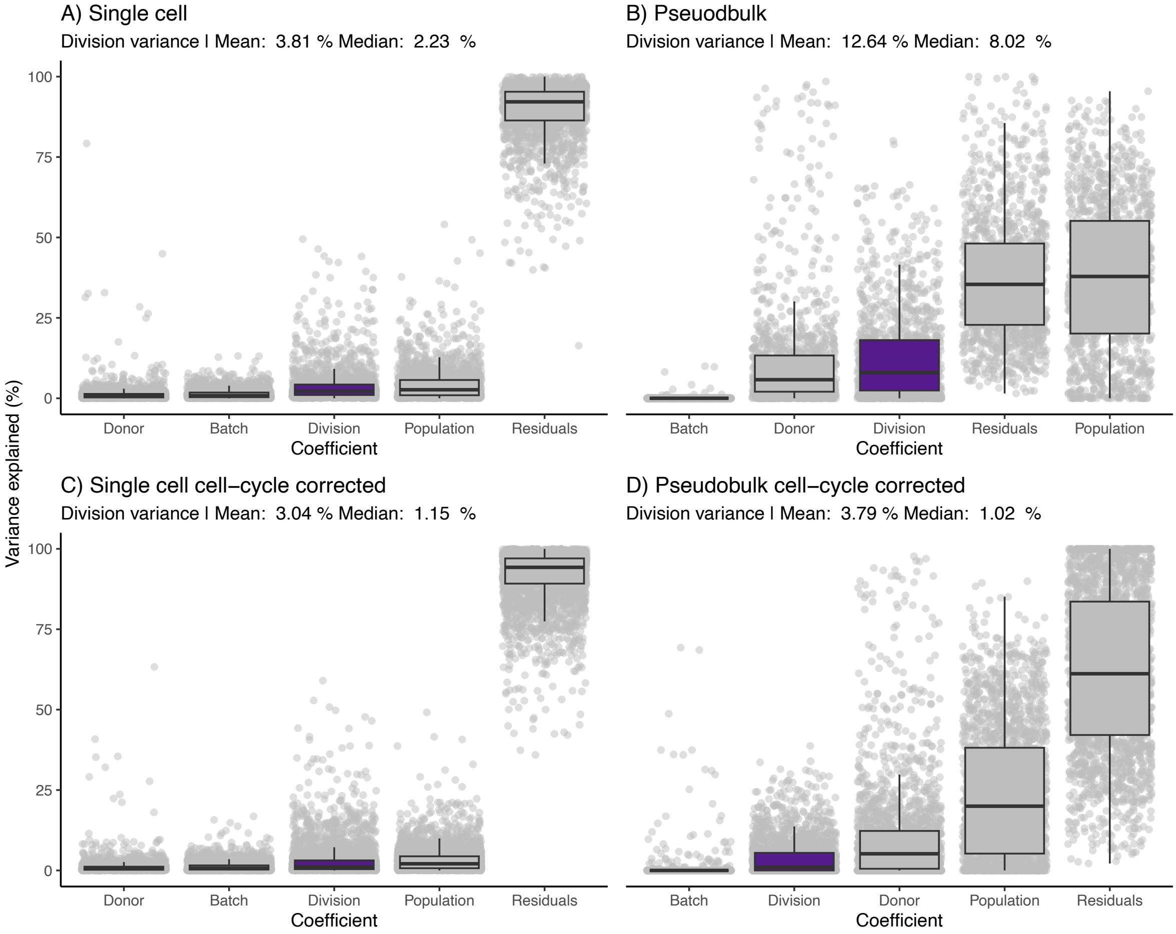


***Supplementary Figure 1 | Gene expression variance explained by division.*** *Variance partitioning analysis showing the proportion of variance explained for the 2,000 most variable genes, calculated from single-cell (A) and pseudobulk (B) log(cp10k+1) normalised expression profiles. C) As A, but showing the variance partition of the residuals after correcting for G2M and S phase scores. D) As B, but showing the variance partition of the residuals after correcting for G2M and S phase scores. G2M and S phase scores were calculated on the pseudobulk sample.*

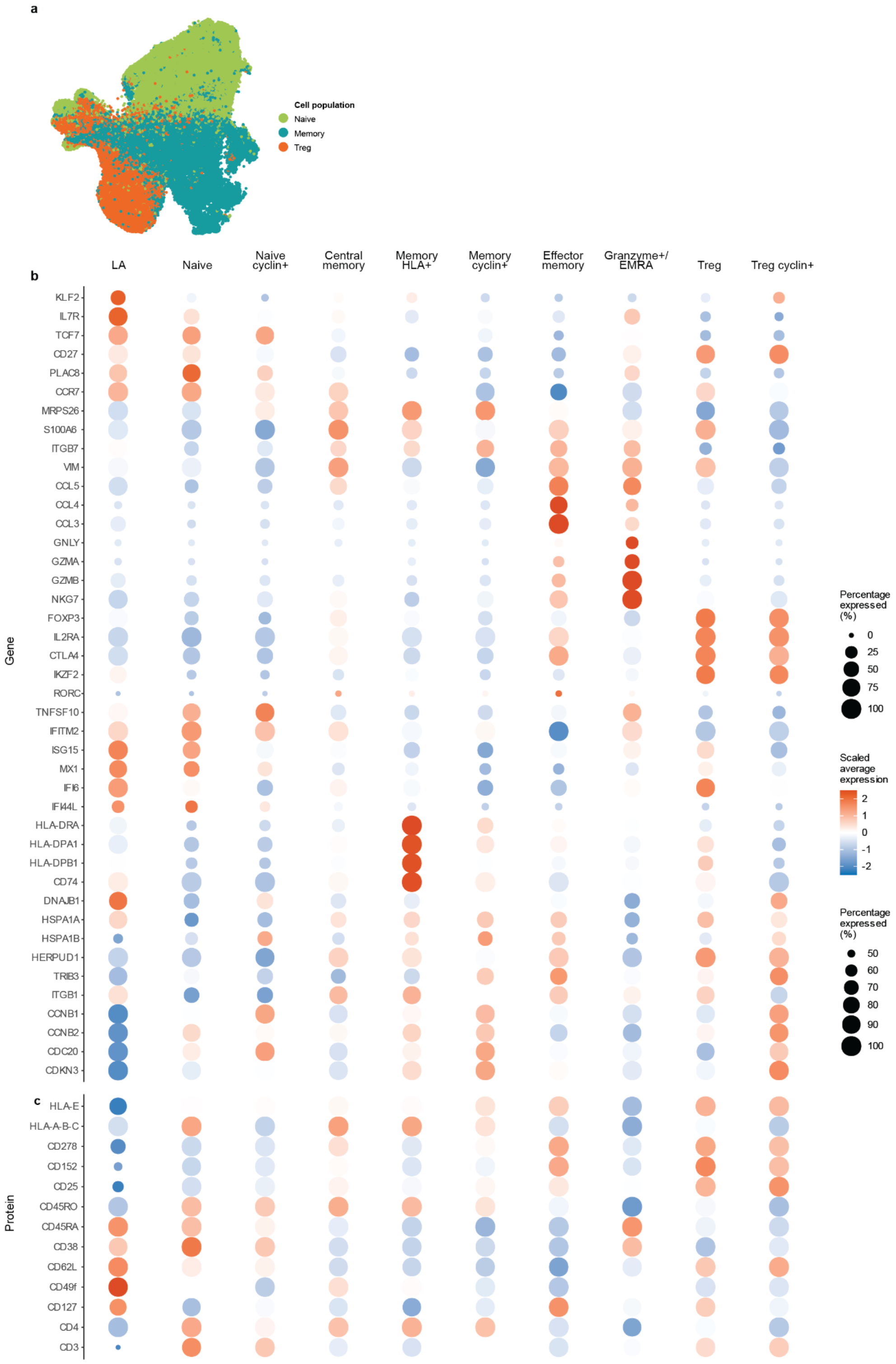


**Supplementary Figure 2 | Cell population and subpopulation annotation.**
a, UMAP embedding of all post-QC cells coloured by population cluster (naive, green; memory, teal; Treg, orange). b-c, Expression of canonical markers used for cluster annotation at the transcriptomic (b) and proteomic (c) levels, corresponding to Figure 1d. Dot size represents the proportion of cells expressing the feature, and colour denotes scaled average expression (red, high; blue, low).

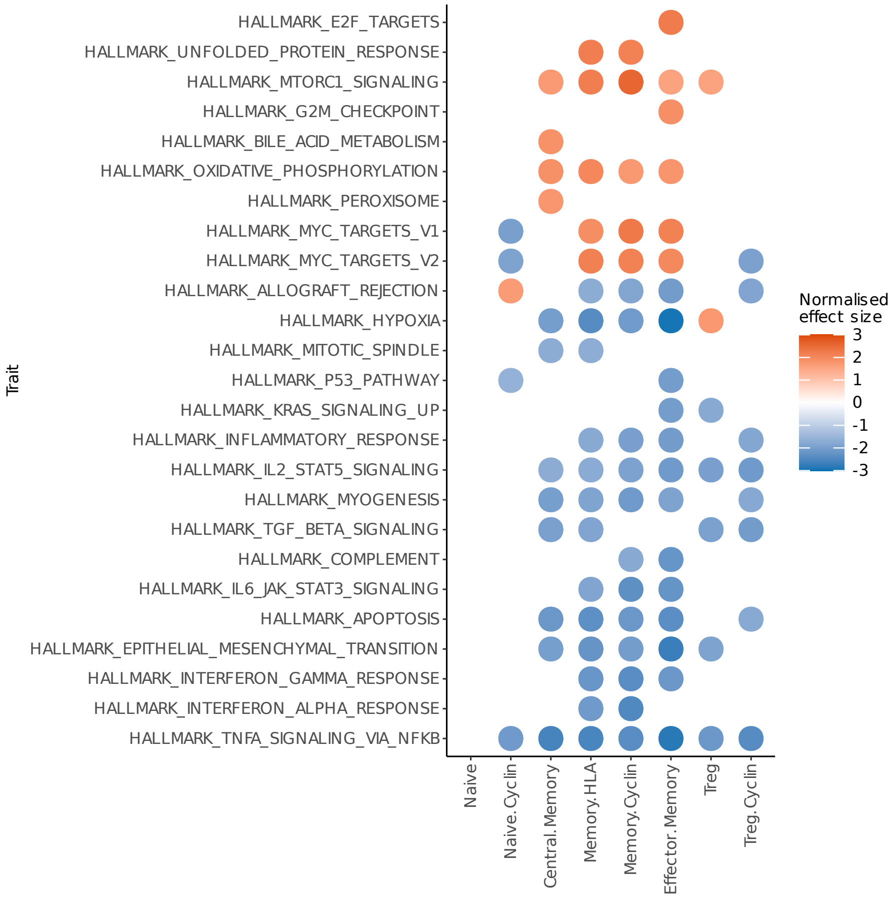


**Supplementary Figure 3 | Gene set enrichment analysis (GSEA) of division-dependent genes.**

Top 10 significantly enriched gene sets (FDR < 0.05) per CD4$^{+}$ T-cell subpopulation from the Hallmark collection (h.all.v2025.1.Hs). Gene sets are coloured by normalised effect size, indicating the direction and magnitude of the division-dependent relationship.

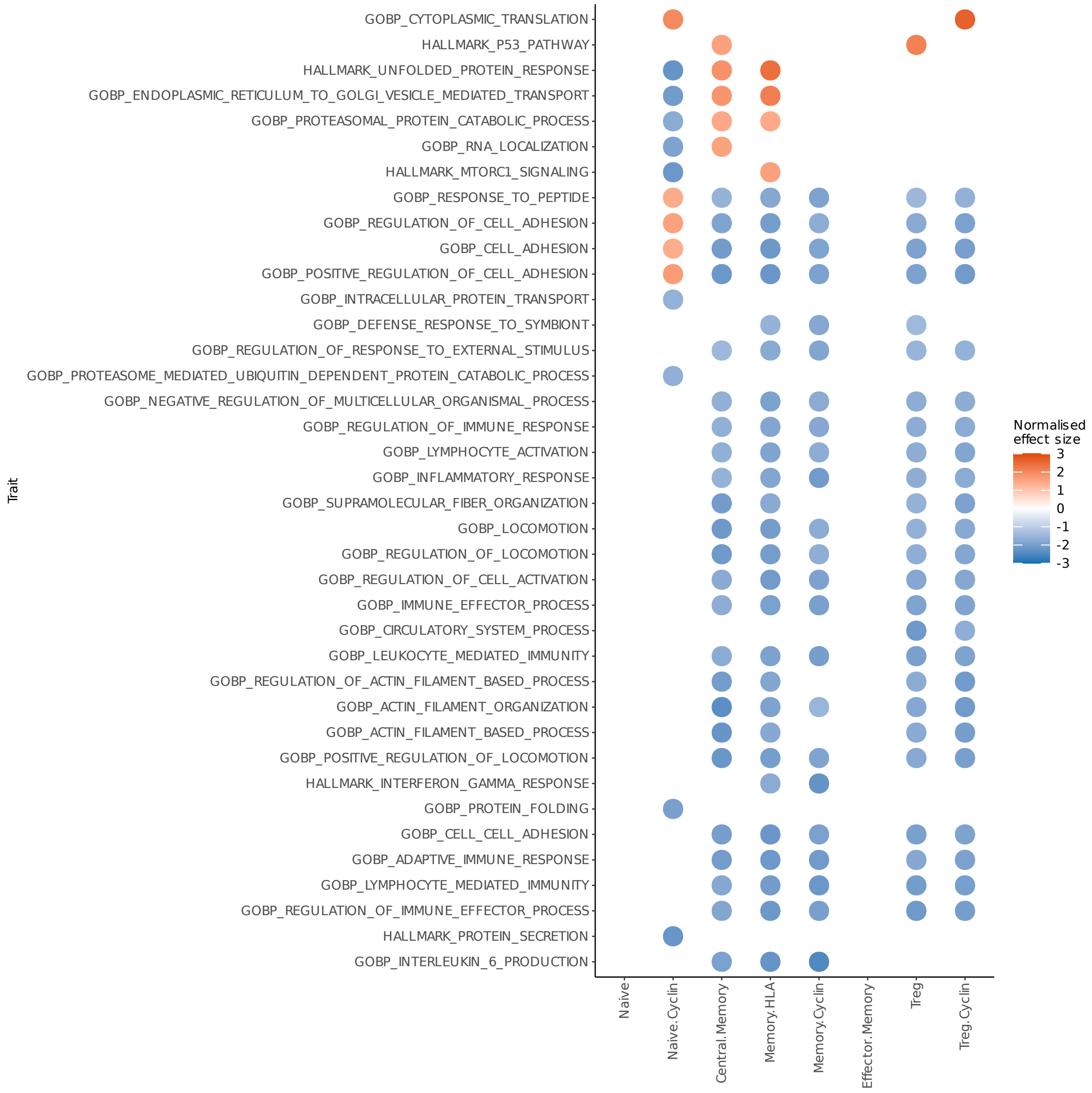


**Supplementary Figure 4 | Gene set enrichment analysis (GSEA) of divergent division-dependent genes.**

Top 10 significantly enriched gene sets (FDR < 0.05) per CD4⁺ T-cell subpopulation from the Hallmark collection (h.all.v2025.1.Hs) and GO Biological Processes (c5.go.bp.v2025.1.Hs). Gene sets are coloured by normalised effect size, indicating the direction and magnitude of the division-dependent relationship.

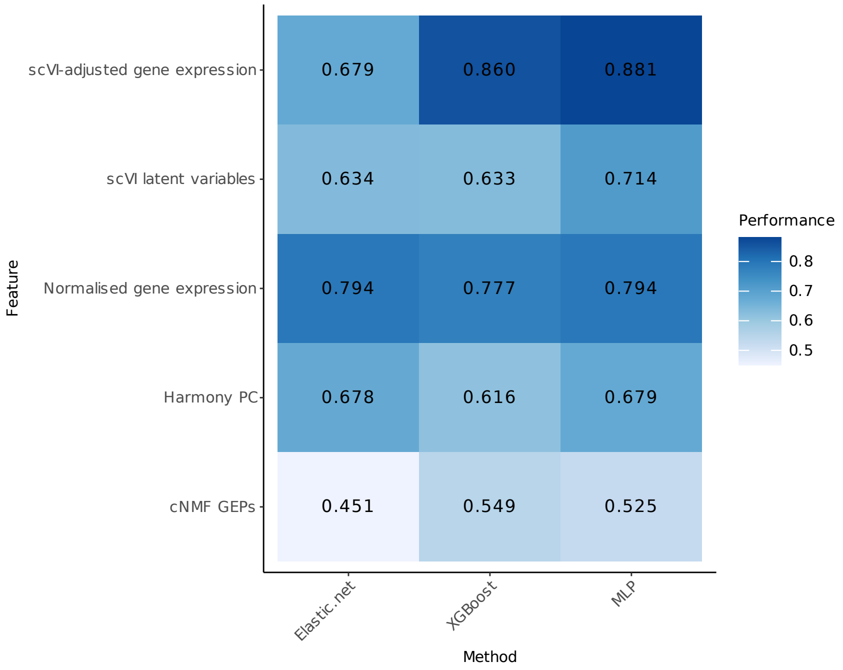


**Supplementary Figure 5 | Optimisation of CellDivider machine learning model.**
Pearson correlation coefficients of predicted and true division states in a held-out test dataset. We evaluated the performance of combinations of five input features and three machine learning methods. Elastic net, Elastic net regression; XGBoost, eXtreme Gradient Boosting; MLP, multi-layer perceptron.

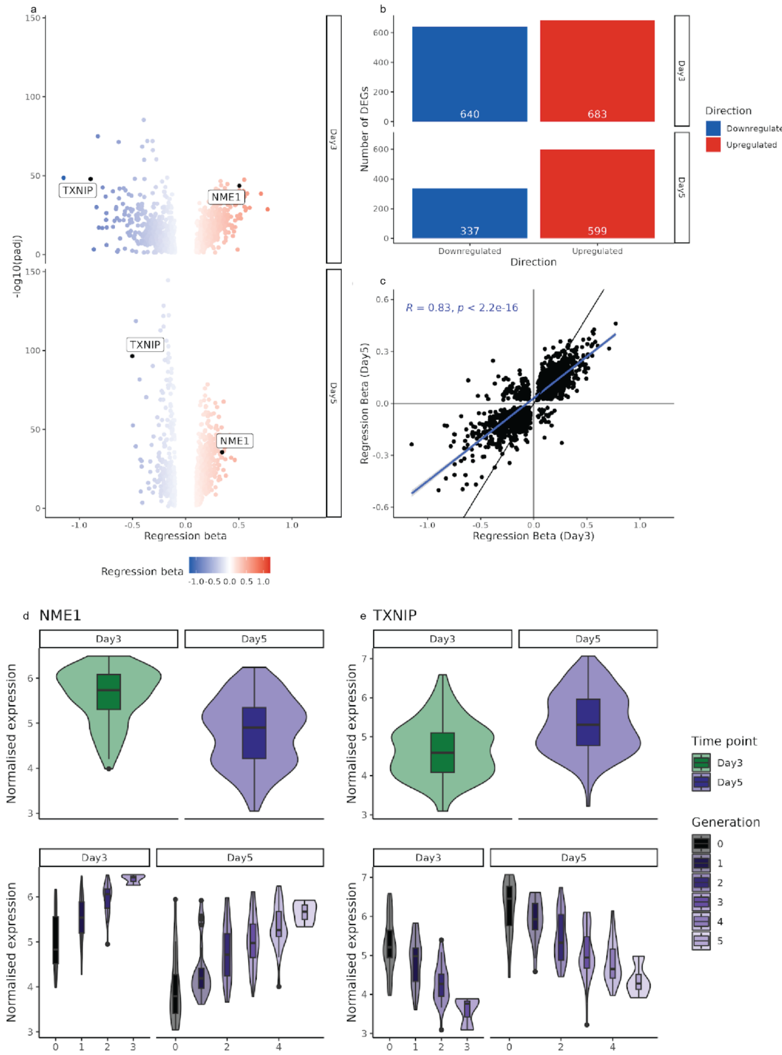


**Supplementary Figure 6 | Highly correlated division-dependent gene expression across Day 3 and Day 5.**

A) Volcano plots showing differential expression results for Day 3 and Day 5, with genes coloured by regression beta (β). Marker genes TXNIP and NME1 are highlighted in black. B) Bar plot showing the number of significantly differentially expressed genes (DEGs) at each time point ($|\beta| > 0.1$, adjusted p-value $< 0.05$), coloured by direction of effect (upregulated vs. downregulated). C) Scatter plot comparing β values of genes detected at both Day 3 (x-axis) and Day 5 (y-axis). The black dashed line indicates the line of equality ($x = y$), while the blue line represents the linear regression fit ($R = 0.83$, $p < 2.2e{-}16$). D–E) Violin and box plots showing normalised metacell expression levels of NME1 (D) and TXNIP (E) at Day 3 and Day 5. For each gene, the top panels display expression values aggregated by time point (i.e., conventional pseudobulk time points), while the lower panels show expression resolved by division state for each time point. These comparisons illustrate that division-resolved profiling provides clearer and more consistent expression patterns than time point-level analysis alone.

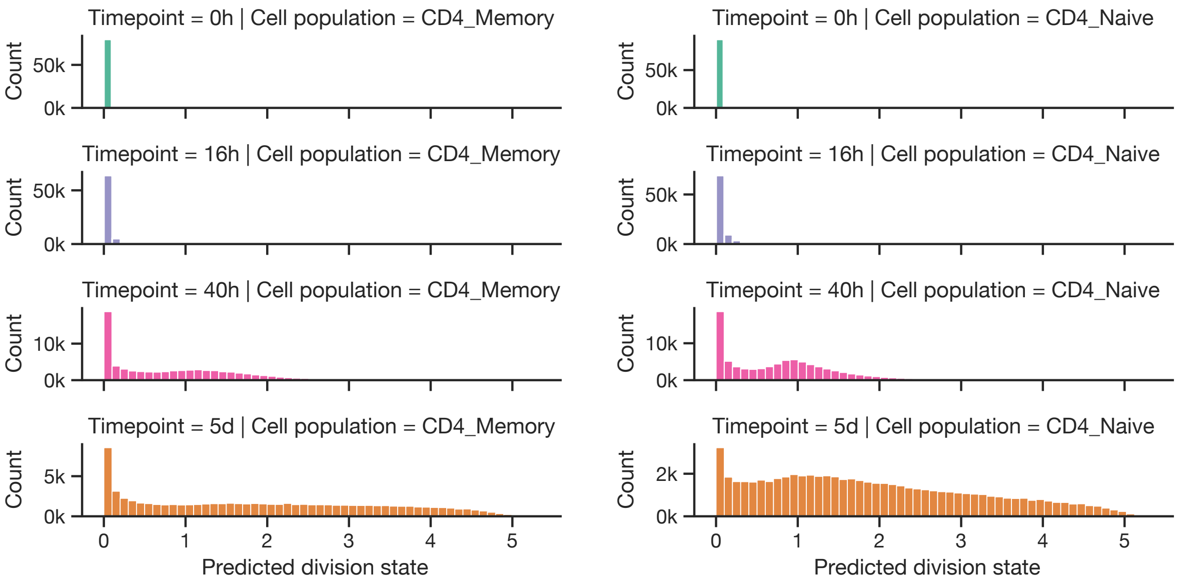


**Supplementary Figure 7 | Predicted division state in the Soskic et al. dataset.** Histograms showing predicted division state distribution, stratified for activation timepoints (0h, 16h, 40h, and 5d) and cell populations (memory and naive CD4$^{+}$ T cells).

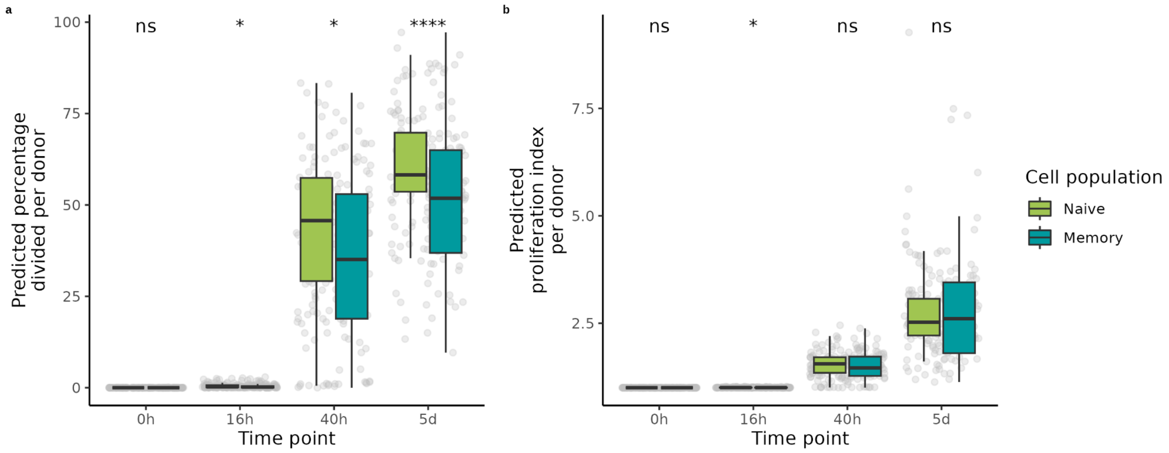


**Supplementary Figure 8 | Predicted proliferation characteristics across cell populations.**

Boxplots show donor-level predicted proliferation metrics for single-cell samples, with boxes coloured by cell population. (a) Percentage of total cells that divided at least once. (b) Proliferation index, defined as the average number of progeny generated per initial cell. Statistical significance between cell populations was assessed using pairwise t-tests within each time point, with P-values adjusted for multiple testing using the Benjamini–Hochberg method. Significance levels are indicated as ns (p > 0.05), * (p <= 0.05), ** (p <= 0.01), *** (p <= 0.001), **** (p <= 0.0001).

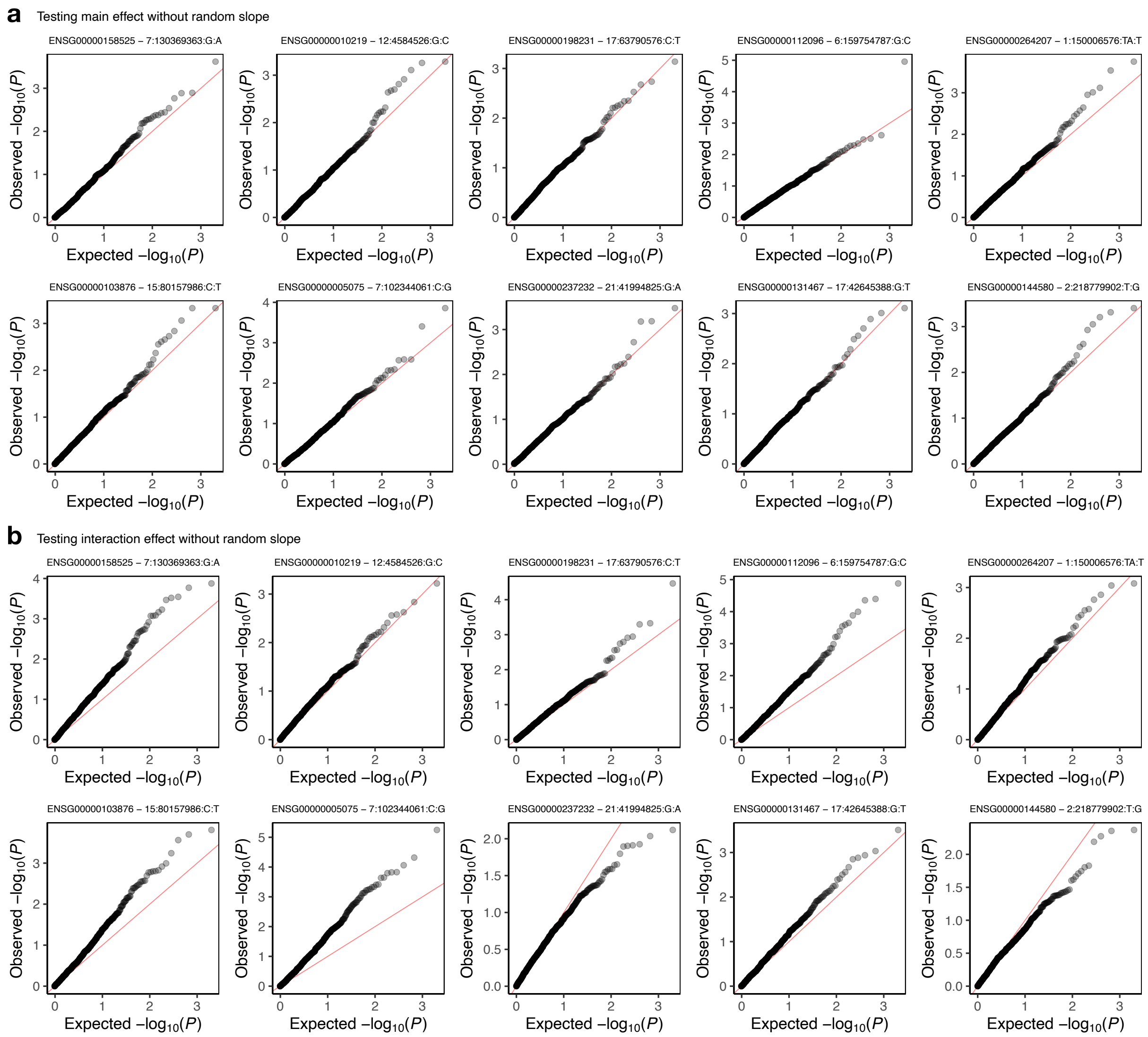


**Supplementary Figure 9 | Permutation experiment of the main and interaction effect testing using the previously proposed NBME model.** Quantile–quantile plots showing statistical calibration for the significance of A) main and B) interaction effects. X-axes indicate uniform p-values and Y-axes indicate observed p-values derived from the likelihood ratio test using 1,000 permuted datasets. Red straight lines indicate identity lines.

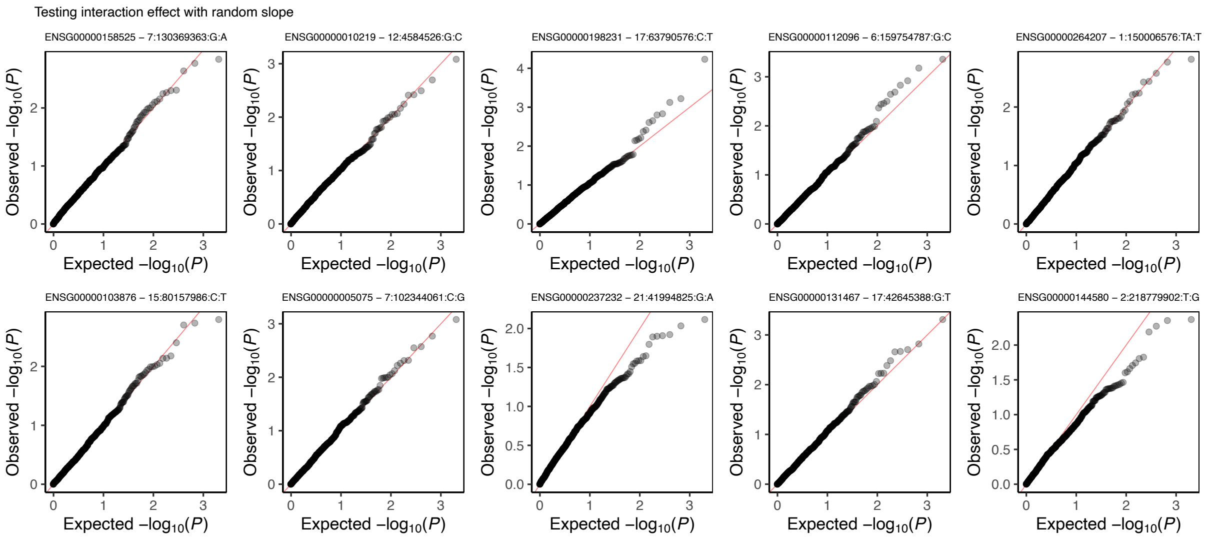


**Supplementary Figure 10 | Permutation experiment of the interaction effect testing using the extended NBME model.**

Quantile–quantile plots showing statistical calibration for the significance of interaction effects. X-axes indicate uniform p-values and Y-axes indicate observed p-values derived from the likelihood ratio test using 1,000 permuted datasets. Red straight lines indicate identity lines.

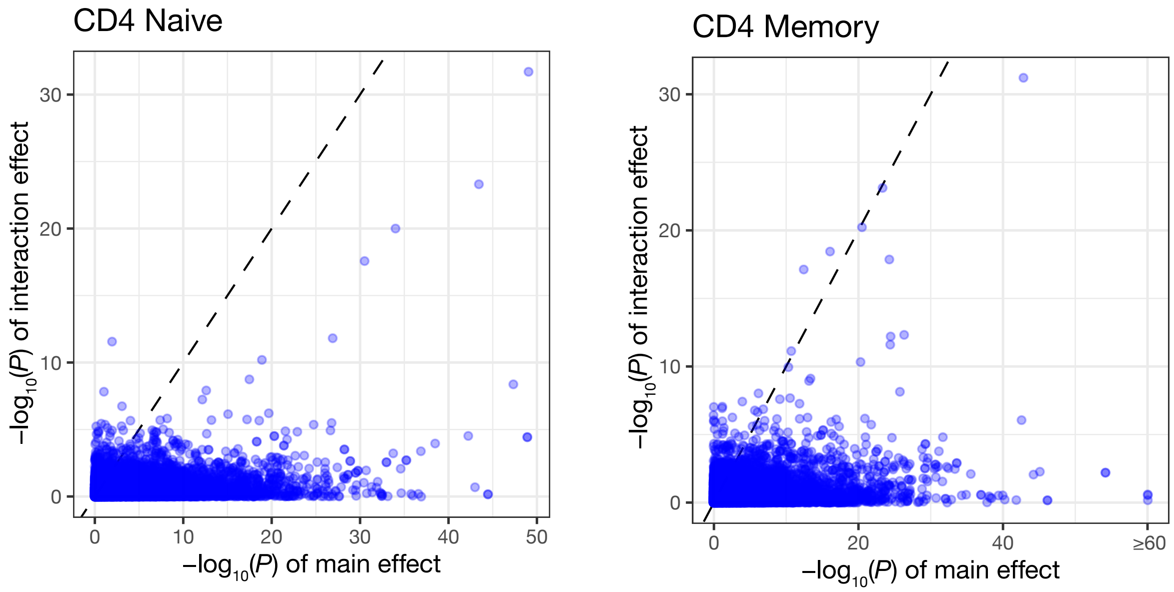


**Supplementary Figure 11 | Comparison of statistical significance of main and interaction eQTL effect**

Scatter plots comparing p-values for main and interaction eQTL effect derived from the likelihood ratio test. Each marker represents an eGene–eVariant pair in Soskic et al. Dashed lines indicate identity lines.

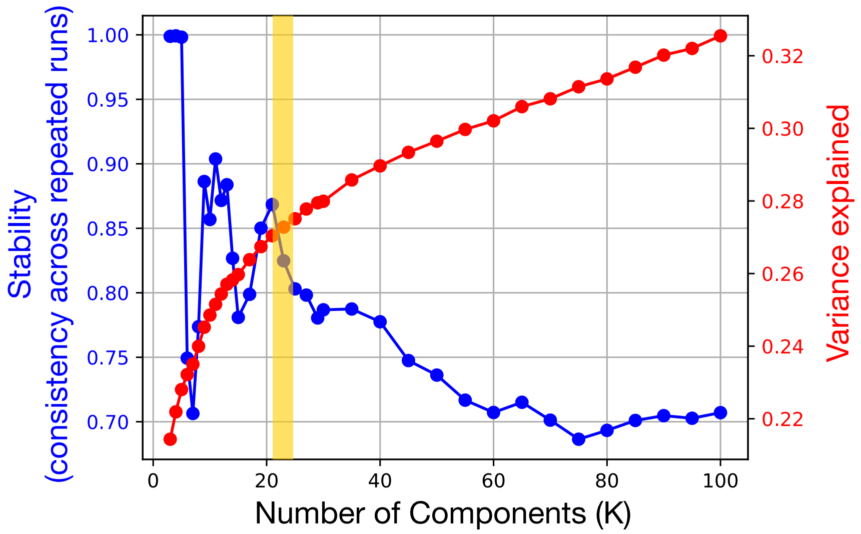


**Supplementary Figure 12 | Optimization of the number of cNMF components.** Optimisation of the choice of the number of components (K). Stability of the components over 100 NMF runs (blue) and variance explained by the defined components (red). The yellow ribbon highlights K = 23, the parameter adopted in our analysis.

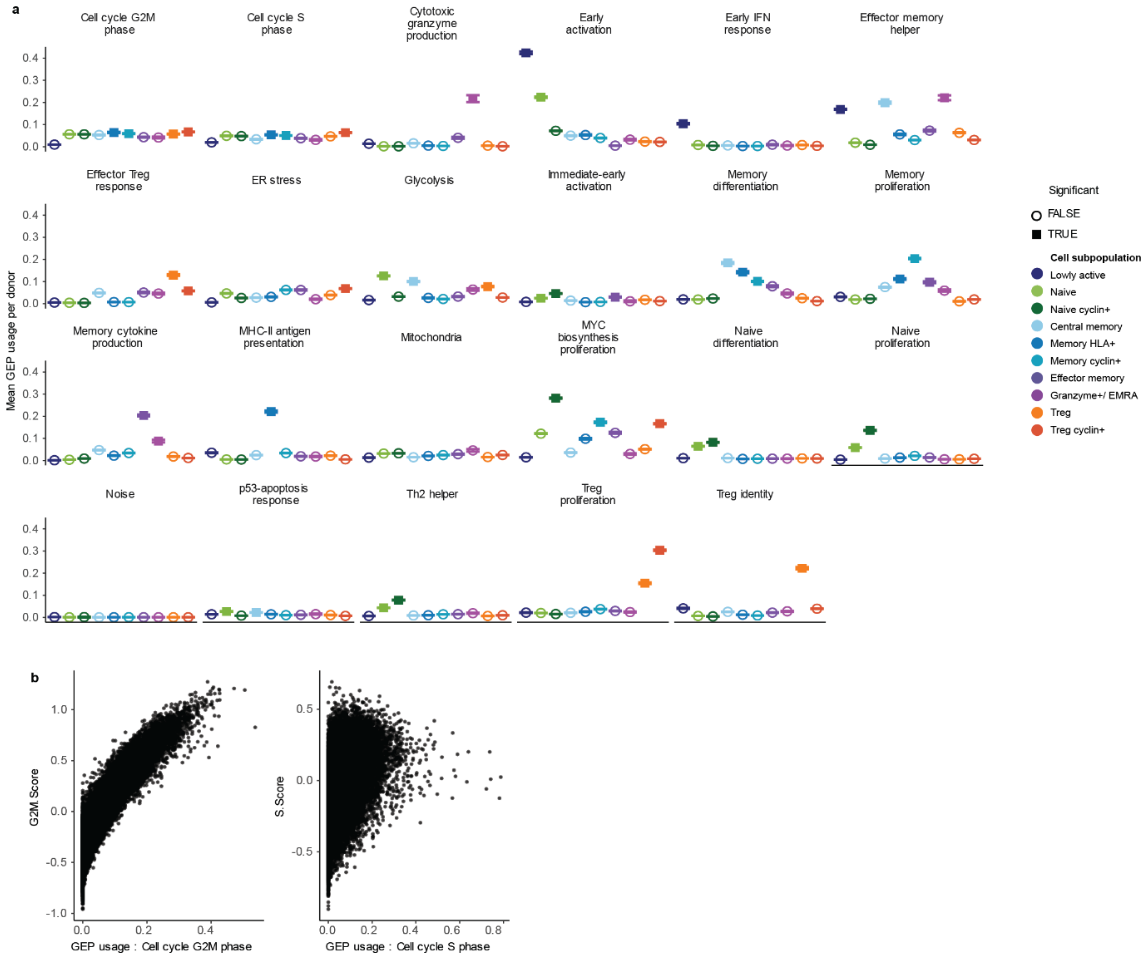


**Supplementary Figure 13 | Cell subpopulation-specific GEP usage and concordance with cell cycle phase scores.**
A) Mean GEP usage across donors for each cell subpopulation (±95% confidence intervals), coloured by cell subpopulation. Solid squares indicate subpopulations with significant GEP usage, as defined in the Methods. B) Correlation between G2/M- and S-phase GEP usage and corresponding G2/M and S phase scores inferred by Seurat CellCycleScoring (**Methods**).

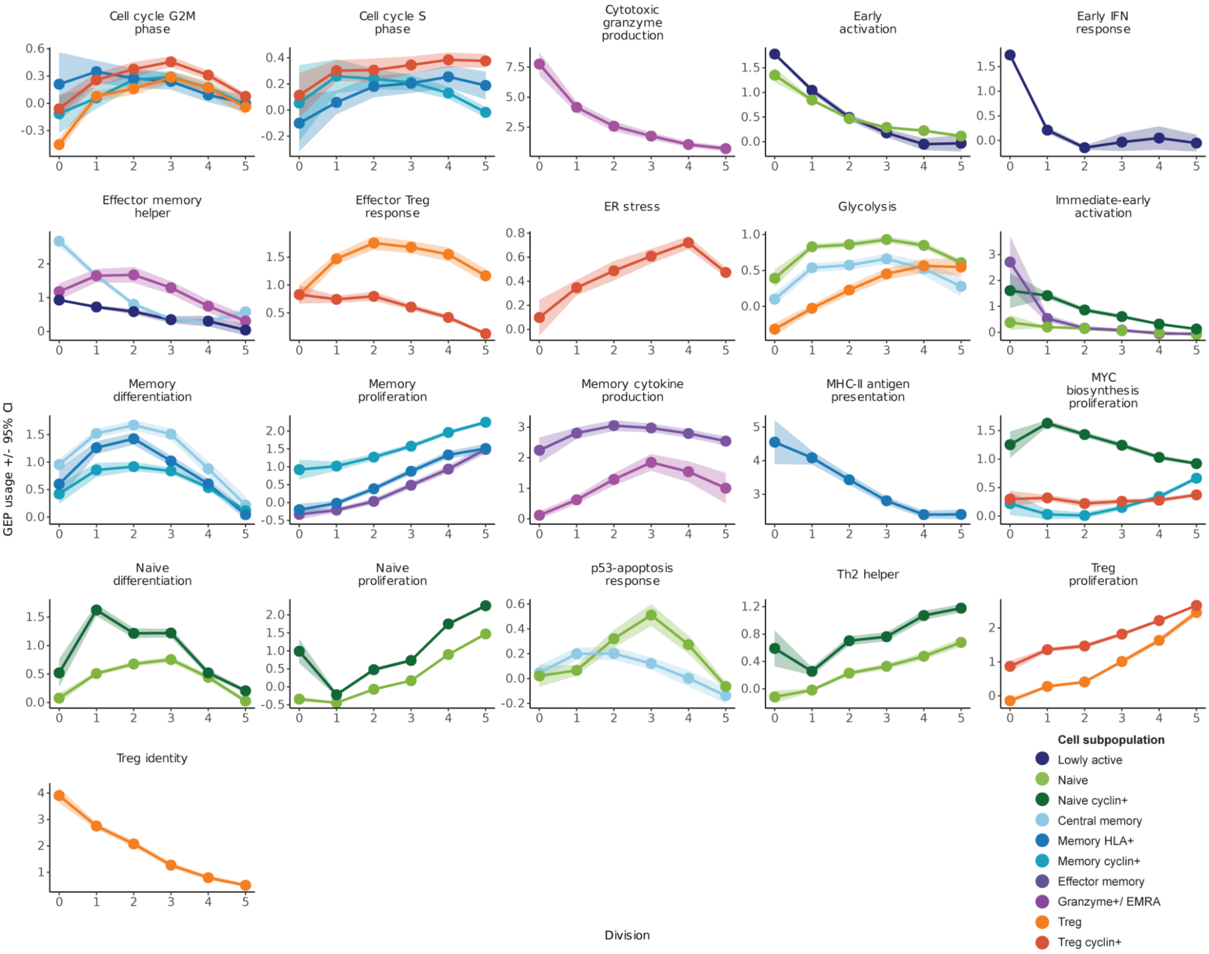


**Supplementary Figure 14 | GEP usage across division states.**
Mean GEP usage across division states for each cell subpopulation (±95% confidence intervals), coloured by cell subpopulation. Only subpopulations with significant GEP usage are shown (**Methods**).

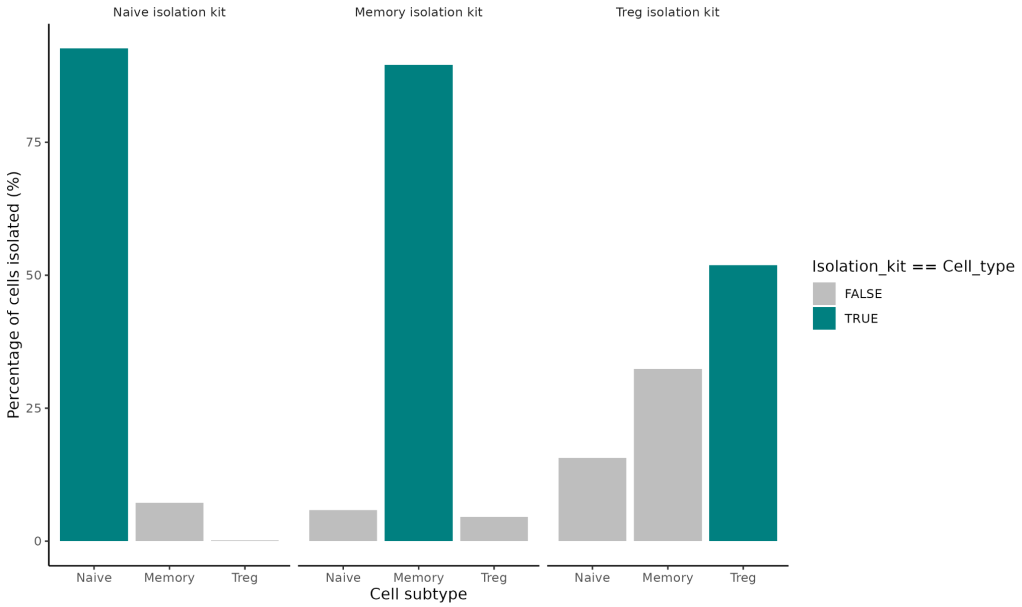


**Supplementary Figure 15 | Contaminant T-cell populations across isolation kits.**
Bar plot showing the number of cells annotated as non-target cell populations by transcriptomic profiling, grouped by isolation kit. Bars are coloured by whether the cell population is a target (teal) or non-target (grey) for the given kit.
